# Heterochiasmy facilitated the establishment of *gsdf* as a novel sex determining gene in Atlantic halibut

**DOI:** 10.1101/2020.11.24.396218

**Authors:** R.B. Edvardsen, O. Wallerman, T. Furmanek, L. Kleppe, P. Jern, A. Wallberg, E. Kjærner-Semb, S. Mæhle, S. K. Olausson, E. Sundström, T. Harboe, R. Mangor-Jensen, M. Møgster, P. Perrichon, B. Norberg, C-J Rubin

**Affiliations:** Institute of Marine Research, P.O. Box 1870, Nordnes, NO-5817, Bergen, Norway; Institute of Marine Research, Austevoll Research Station, NO-5392, Storebø, Norway; Department of Medical Biochemistry and Microbiology, Uppsala University, 75123 Uppsala, Sweden

**Keywords:** aquaculture, nucleotide diversity, transposable elements, genome assembly, meiotic recombination, sex chromosome

## Abstract

Atlantic Halibut (Hippoglossus hippoglossus) has a X/Y genetic sex determination system, but the sex determining factor is not known. We produced a high-quality genome assembly and identified parts of chromosome 13 as the Y chromosome due to sequence divergence between sexes and segregation of sex genotypes in pedigrees. Linkage analysis revealed that all chromosomes exhibit heterochiasmy, i.e. male- and female restricted meiotic recombination intervals (MRR/FRR). We show that FRR/MRR intervals differ in nucleotide diversity and repeat class content and that this is true also for other Pleuronectidae species. We further show that remnants of a Gypsy-like transposable element insertion on chr13 promotes early male specific expression of gonadal somatic cell derived factor (gsdf). Less than 4 MYA, this male-determining element evolved on an autosomal FRR segment featuring pre-existing male meiotic recombination barriers, thereby creating a Y chromosome. We propose that heterochiasmy may facilitate the evolution of genetic sex determination systems.

## Introduction

The Atlantic halibut (*Hippoglossus hippoglossus*) is the largest teleost species in the North Atlantic and can reach more than 50 years of age. In recent years, the demand for Atlantic halibut has been larger than the supply of wild fish. In some areas the stocks have been defined as vulnerable and sensitive to overfishing, due to relatively slow growth and high age at maturity. In combination with a high market price, this has led to an increased interest in halibut aquaculture. Female halibuts can weigh more than 350 kg and become more than 3.6 m long, while males rarely weigh more than 50 kg. A problem encountered in Atlantic halibut aquaculture is that males mature earlier and at a smaller size than females^1^, and efficient elimination of male halibut in production of market fish would therefore constitute a particularly great advantage in halibut compared to other aquaculture fish species. Atlantic halibut has an XX/XY, male heterogametic, sex determination system^2^. The most common and practical method to achieve production of all female populations is sex reversal of F1 female juveniles, to create phenotypic and functional males with an XX karyotype^3,4^. The offspring of these sex-reversed genetic females will be 100% female, but identifying and separating XX males from XY males involves raising offspring sibling groups until their sex can be visually determined by histology or microscopy of gonads^4,5^, a time- and resource-consuming process that can take up to four years. A reliable and practical method for determination of genetic sex based on identification of Y chromosome markers is instrumental for the development of efficient protocols for all-female production.

In contrast to the highly conserved mammalian and avian sex determination systems, fish generally display much more plasticity and turnover when it comes to sex chromosomes and sex determination systems, with many already described systems being young in comparison to their mammalian and avian counterparts^6^. Furthermore, teleosts display a rich diversity of sex-determining mechanisms, both environmental and genetic^7^. As for genetic control of sex, both XX/XY and ZZ/ZW systems have been described in teleosts, sometimes with closely related species having differing systems, such as for sticklebacks^8^, tilapias^9^, swordtail fish^10^ and mosquitofish^11^. Righteye flounders (*Pleuronectidae*) have variable genetic sex determinations systems with both XX/XY and ZZ/ZW systems represented^12^. The Pacific Halibut (*Hippoglossus stenolepis*), which diverged from the Atlantic halibut about 3.7 million years ago^2^, has a ZZ/ZW system^13^. Several master sex determining (MSD) genes have been identified in teleosts, most of which belong to one of three protein families (DMRT, SOX, and TGF-ß and its signaling pathway)^14^, and these have originated either from sub- or neo-functionalization of duplicated genes, or by allelic diversification^6^. Exceptions to these protein families are found for salmonids, where *sdy* is the MSD gene^15^ and recently the *breast cancer anti-estrogen resistance protein 1* (*BCAR1*) and *bone morphogenetic protein receptor type-1B* (*bmpr1b*) have been reported as MSD genes in catfish (*Ictalurus punctatus*)^16^ and Atlantic herring (*Clupea harengus*)^17^, respectively. Thus, a wide range of systems have evolved for genetic control of sex determination in fish and additional complexity is added by environmental variables contributing to sex determination in some species.

Single molecule sequencing of long reads has transformed the way the scientific community produces genome assemblies^18^. In relation to short-read sequencing technologies, methods capable of sequencing long reads can greatly increase the contiguity of genome assemblies due to long reads bridging repetitive parts of the genome. In this project we used an Oxford Nanopore Technologies (ONT) MinION device to sequence long native DNA molecules from a single male Atlantic halibut and used the assembled contigs in conjunction with Chromatin Conformation Capture sequencing (HiC) data to produce a chromosome-scale scaffolded genome assembly. To identify and characterize the sex chromosome we mapped short read data from male and female DNA pools onto the assembly to screen for divergent genetic markers between the sexes. The differing systems within the *hippoglossus* genus indicates that at least one of Atlantic- and Pacific halibut must have evolved a novel sex determination system after their 3.7 MYA split and that the mechanism for genetic sex determination in Atlantic halibut may be recent on an evolutionary time scale. Thus, our high-quality Atlantic halibut genome assembly offers great potential to study the emergence of a young genetic sex determination system and evolution of sex chromosomes.

## Results

### Genome assembly and annotation

We used nanopore sequencing in combination with Illumina sequencing of 10X Genomics Chromium-, paired-end- and HiC libraries to generate a chromosome-scale assembly of the Atlantic halibut. The final scaffolded assembly (IMR_hipHip.v1) had a scaffold N50 of 27 Mb (Supplementary Fig. 1a,b) and this assembly was evaluated using BUSCO using 4584 genes in *Actinopterygii* odb9 data set, annotating 95.4% of analyzed genes as “Complete” in the genome assembly. The assembly is organized into 24 larger scaffolds containing 97.6% of the assembled sequence (Supplementary Fig. 2), corresponding to the chromosomes inferred from karyotyping^19^ and linkage analysis^20^. A number of smaller contigs and scaffolds, together comprising only 2.4 % of the assembled sequence were left as unplaced. We named the scaffolds by mapping microsatellite primer sequences from a published genetic linkage map for Atlantic halibut^20^ to IMR_hipHip.v1. Genome to genome alignment of IMR_hipHip.v1 to a recently finished reference genome of the Pacific halibut (NCBI accession GCF_013339905.1) revealed a largely co-linear genome organization, with few large-scale disagreements (Supplementary Fig. 1c) as expected due to the recent split of the two species. To generate RNA-seq data for genome annotation and to obtain transcriptomic data describing the early stages of development, we used samples from different time points throughout embryonic development from newly fertilized to 54 days post fertilization (dpf). For the first four stages (1 hour post fertilization, 5 dpf, 8 dpf and 12 dpf), four pools of eight individuals were collected and subjected to RNA extraction and Illumina HiSeq mRNA-sequencing. For the 28 dpf, 43 dpf and 54 dpf, four individual fish were sampled and sequenced for each stage. Genome annotation based on the RNA-sequencing identified 18,314 genes, of which 18,092 (98.8%) significantly matched with proteins in public databases.

### Identification of the Atlantic halibut sex chromosome

In order to identify and characterize the sex chromosomes we sampled and extracted and pooled DNA from ten full siblings of each sex and sequenced these using Illumina HiSeq sequencing. The sequencing data was mapped against the IMR_hipHip.v1 assembly and the sequence divergence between the male and female pools were determined for non-overlapping windows of 100 SNPs along each chromosome using χ^2^-tests comparing male- and female allele frequencies. This screen revealed chromosome 13 as the most diverged between males and females (Fig. 1a), in agreement with a previously published linkage study using RAD-sequencing^21^. It should be noted that the haploid nature of the IMR_hipHip.v1 assembly makes it possible that chr13 in this assembly is a hybrid between an X and a Y, which is what we find when comparing the pool-seq genotypes of male vs. female divergent SNPs. The divergence between phenotypic males and females on chr13 is not uniform, but overall, gradually declines along the length of the chromosome, with the end being no more divergent than autosomes (Fig 1b and c).

**Fig. 1:**
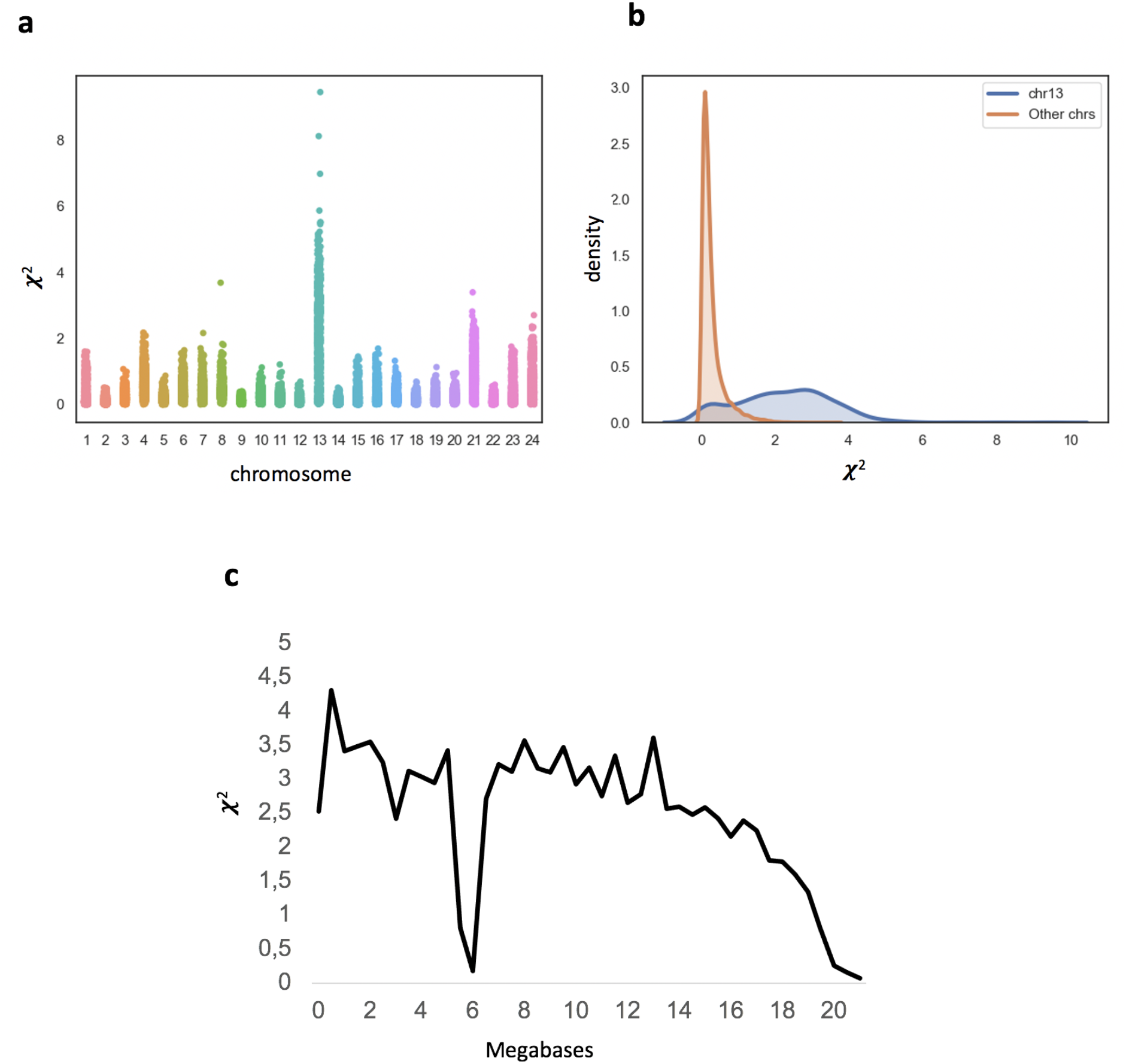
Genetic differentiation between phenotypic males and females indicates chr13 as the Y-chromosome. **a** Average *χ*^2^-values for 100 SNPs along each assembled chromosome in the Atlantic halibut genome. **b** Distribution of average *χ*^2^-values for chr13 compared to the rest of the genome. **c** Average ***χ***2-values for 500 kb windows along chr13.

### Identification of the Atlantic halibut sex determining gene

RNA-seq data was generated from 12 individuals covering three developmental stages with the assumption that both genetic sexes would be represented. As we were unable to infer phenotypic sex for these samples, an unbiased screen could involve searching for the most variably expressed genes among the 12 individuals, but after conducting a Principal Component Analysis we concluded that the samples clustered strongly by development stage (Supplementary Fig. 3). An alternative strategy could involve screening for variable gene expression between known phenotypic sexes during gonadal differentiation. As known phenotypic sex was unavailable, we determined the genetic sex of the RNA-seq samples (XY or XX). We first mapped the DNA-seq pools to the genome and extracted SNPs that had approximately 50% allele frequency in the male pool-seq while being fixed for one allele in the female pool. We next mapped the RNA-seq samples to the genome, extracted RNA-seq alleles at the extracted divergent positions and performed hierarchical clustering (Supplementary Fig. 4) showing clearly that 7 samples were genetically male (XY) and 5 were genetically female (XX). We then produced a Volcano plot showing the global genetic male vs. female expression ratio plotted against the statistical significance of a differential expression (DE) between sexes (Fig. 2a). The most significantly DE gene was *gsdf*, previously shown to be the male-determining MSD gene in Luzon ricefish and sablefish^22,23^. Furthermore, among genes previously identified as MSD genes in other fish species^14^, only *gsdf* resides on chr13 in Atlantic halibut (chr13:8506700-8510213). Among 12 RNA-seq individuals the 5 XX fish had almost no *gsdf* expression, while the 7 XY fish all had high expression, consistent with Gsdf being a male determining factor. To verify sex specific expression of *gsdf* during development and to obtain better temporal resolution of expression levels, we performed quantitative PCR (qPCR) on PCR sex-determined samples (Fig. 2b and c). RNA was extracted from individual embryos from 11 development stages, ranging from 1 hour post fertilization (hpf) to 13 dpf. A PCR assay based on SNPs in the 3’UTR of *brx* (chr13:9125004-9125007) was used to determine genetic sex of samples from 48 hpf and onwards (Supplementary Fig. 5). The qPCR results show that the expression of *gsdf* starts in males already at 48 hpf, and from 7 dpf onwards there is a clear increase in expression compared to the earlier stages. For one individual sampled at 6 dpf determined as a genetic male, we could not detect any expression, possibly due to non-genetic factors having the chance to overrule the genotype in the determination of phenotypic sex in the species. *in silico* searches for conserved genes located up- and downstream of *gsdf* confirmed the *gsdf* identity. (Supplementary Fig. 6).

**Fig. 2:**
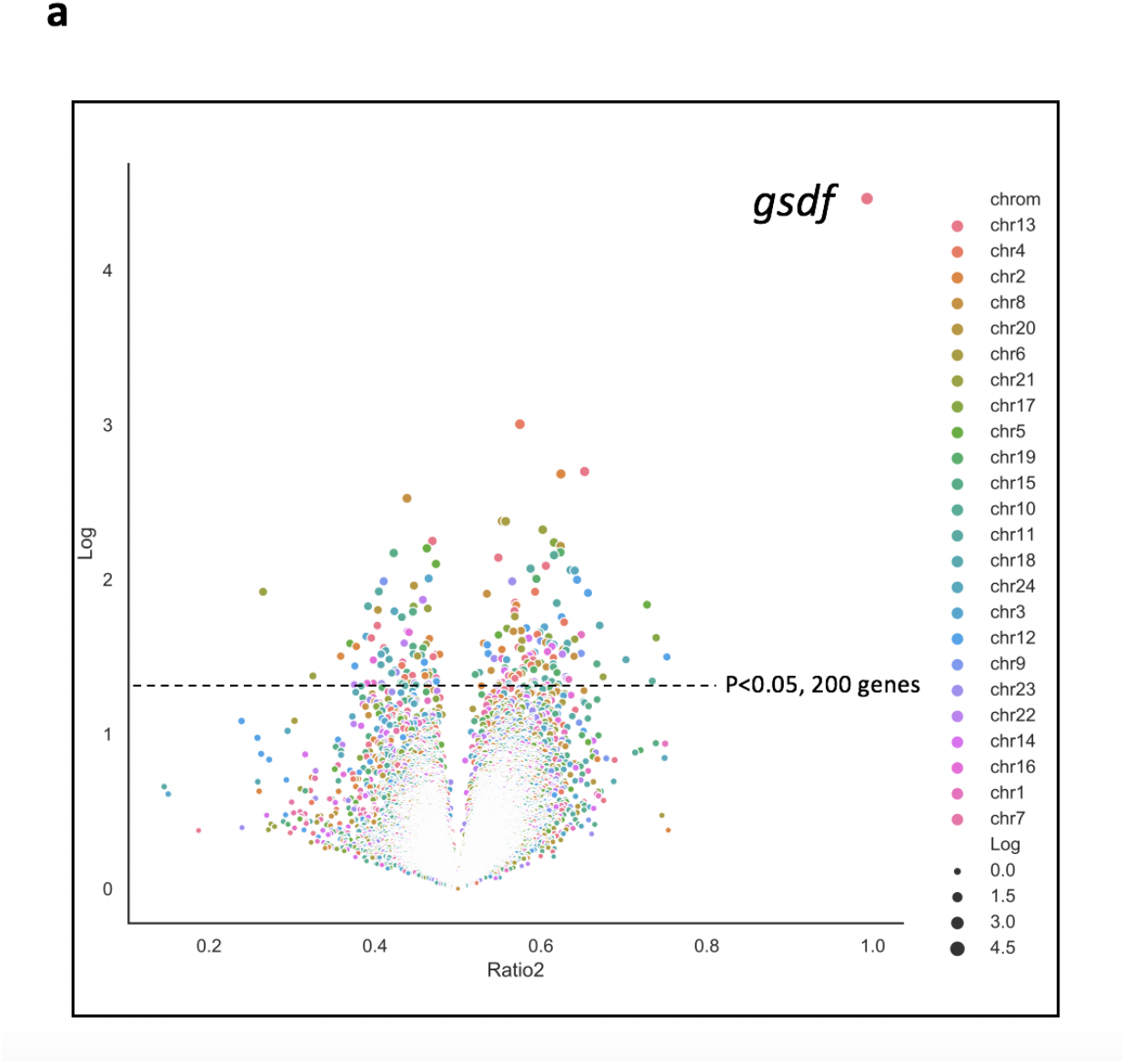
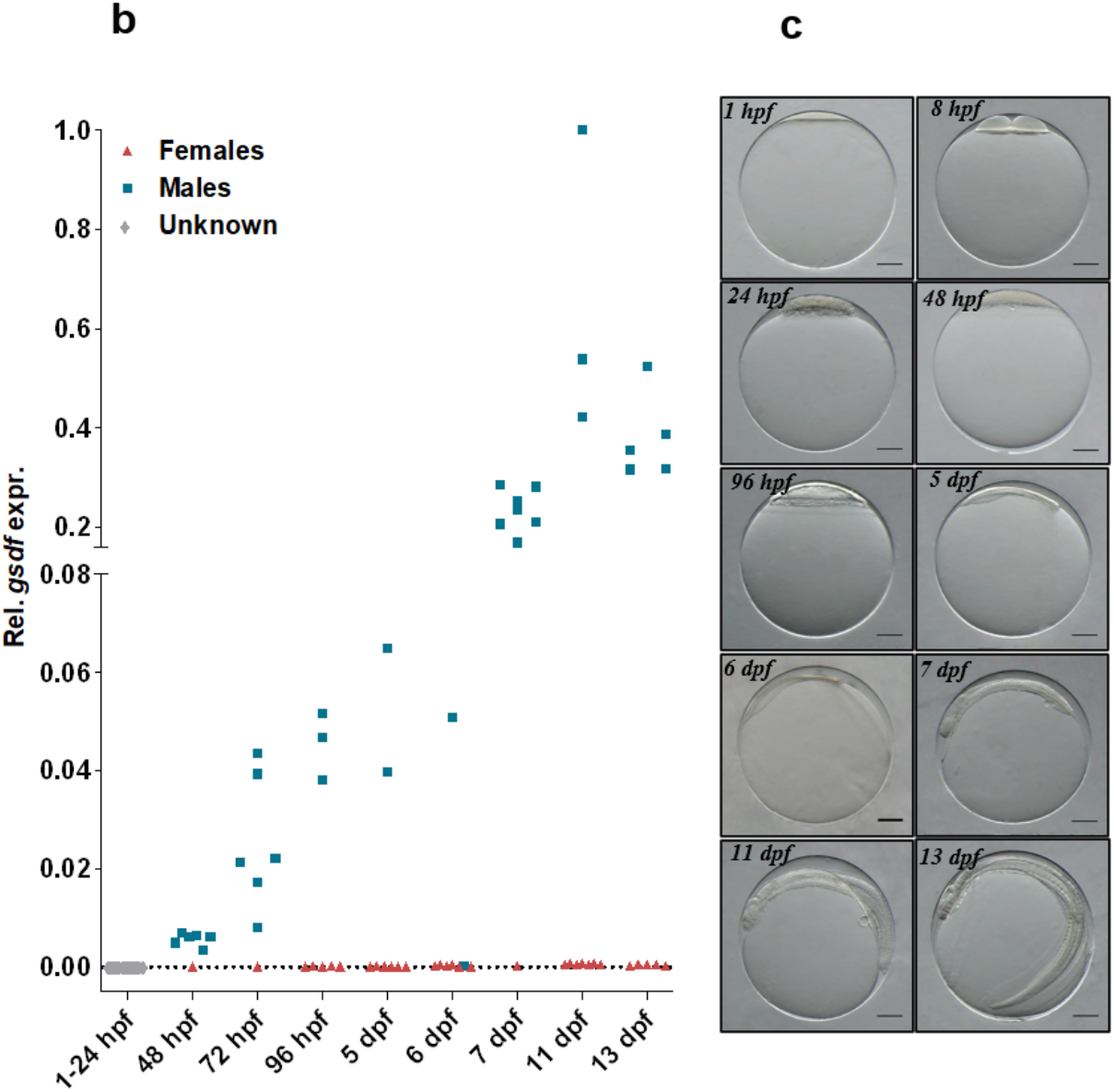
Identification of *gsdf* as the Atlantic halibut sex determining gene. **a** Volcano plot representation of differential expression (DE) between individuals stratified by chr13 genotyping-based sex assignment. Dot colors indicate the chromosome from which genes are expressed and dot sizes were made proportional to the significance of DE (higher significance=larger dots). The most highly differentially expressed gene, *gsdf*, is indicated in the upper right corner. **b** qPCR result showing the sex specific *gsdf* expression (relative to *gtf3c6*) across developmental stages. **c** Pictures of fish representing the developmental stages sampled for gene expression analyses; 1 hpf to 13 dpf.

### Most of the Atlantic halibut genome is subject to male or female restricted meiotic recombination

In order to validate chr13 as the Atlantic halibut sex chromosome we downloaded the pedigree RAD-sequencing data previously used to assign linkage group 13 as X/Y^21^, aligned the data to our assembly IMR_hipHip.v1, called SNPs and investigated genotypes by sex phenotype in the pedigrees. The pedigree analysis verified our divergence-based assignment of IMR_hipHip.v1 chr13 as the sex chromosome and 90/94 F1 individuals were in agreement with regard to the association between phenotypic sex and chr13 genotype. A similar observation was made for the embryonal qPCR analysis where one out of 34 genetic males (*brx*-assay) had no *gsdf* expression. The reason for the non-perfect association between genetic- and phenotypic sex could be erroneous annotation of the downloaded RAD-seq dataset or a logical consequence of non-perfect penetrance of young genetic sex determination systems. In our genome assembly-guided linkage analysis, we detected suppression of meiotic recombination along a large part of chr13 in male germ cells as would be expected in an X/Y system. Surprisingly, the majority of the sex chromosome pair exhibited strict male- or female restricted meiotic recombination (MRR/FRR) (Fig. 3a, Supplementary Fig. 7a). The male- and female genetic maps both sum up to 49 cM but the recombining intervals are nonoverlapping. None of the 90 F1 offspring used for analysis carried a male meiotic recombination in the first ∼15 Mb, while all detected female meiotic recombination events on chr13 occurred in this region. Thus, the first ∼15 Mb of chr13 behaves like a X/Y chromosome, with high male vs. female divergence (Fig. 1c) and nonexistent meiotic recombination in males. The later part of chr13 (∼15 Mb until chromosome end) only recombines in males (X/Y recombination) and has lower male vs. female divergence, suggesting that sequence diversity in this part of the chromosome is shared between X and Y. Even more surprising than the meiotic recombination barrier observed for the of chr13 in females was the observation that all chromosomes clustered into more or less strict MRR/FRR regions (Fig. 3b, Supplementary Table 1 and Supplementary Fig. 7a). We limited our analysis to the placed chromosome part of the genome (chromosomes 1-24), spanning 596 Mb out of the total 611 Mb, and defined FRRs and MRRs according to our classification outlined in Supplementary Table 2. Genome-wide, this annotation defined 407 Mb (68% of the sequence) as FRRs and 154 Mb as MRRs (26% of the sequence) and only ∼6% of the sequence fell outside of this classification, i.e. either both sexes or neither sex showed meiotic recombination.

**Fig. 3:**
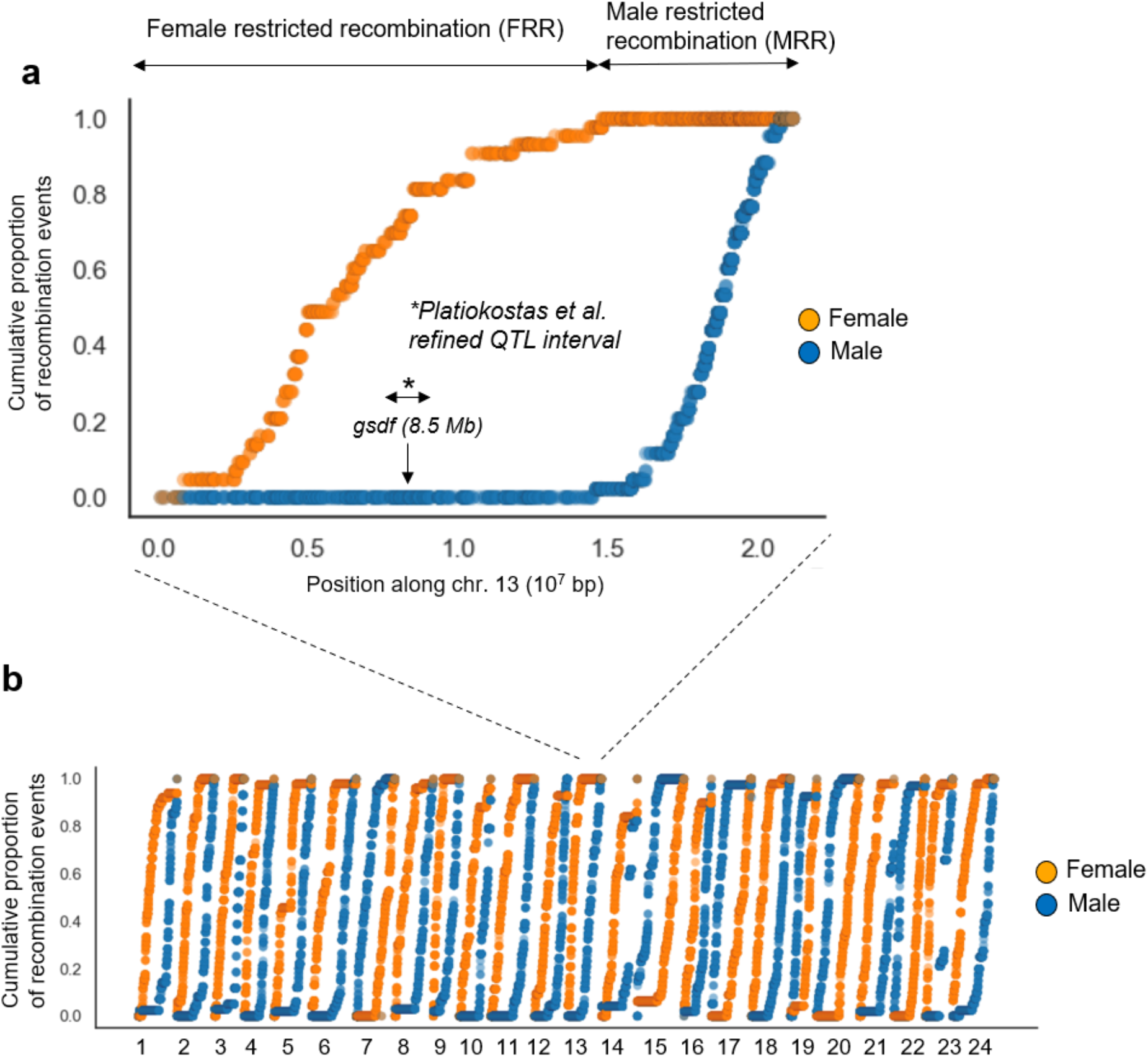
Cumulative proportion of all recombination events occurring during male and female meiosis in a two-pedigree intercross. **a** chr13 with *gsdf* location indicated. *Mapping interval of markers most highly associated with sex in a previous study^21^. **b** Male and female linkage maps differ strongly. All 24 chromosomes exhibit clear sex restricted recombination intervals.

### Nucleotide diversity and repeat content is correlated with sex-restricted recombination

We investigated what distinguishes an MRR from an FRR and concluded that for all chromosomes except chr9 MRRs are much smaller than FRRs while retaining approximately the same number of recombination events (Supplementary Fig. 7b). Furthermore, MRRs, again with the exception of chr9, exhibit higher nucleotide diversity (*π*) than their FRR counterparts (Fig. 4a). Recombination rate is correlated with *π* (Supplementary Fig. 8) and chr13 is an outlier with the largest MRR over FRR *π* ratio among all chromosomes, presumably due to the limited effective population size of its FRR, the herein identified Y-chromosome. Typically, MRRs occupy one end of a chromosome and FRRs start from the other end. Atlantic halibut has been shown to have subtelomeric or acrocentric centromeres^19^ and for 20 out of 21 chromosomes where we could infer centromere placement by mapping centromeric markers^20^ to IMR_Hiphip.v1, the female end coincided with the centromere (Supplementary Table 3). We next sought to determine whether meiotic recombination is sex restricted also in species closely related to Atlantic halibut but could not find any publicly available pedigree datasets for other Righteye flounders. Since Atlantic halibut *π* was found to be higher for MRRs than FRRs, we decided to investigate whether *π* is correlated with MRR/FRR classification also in other Righteye flounder species. We could not find any publicly available genome sequencing data for Pacific halibut and instead used RNA-sequencing data to estimate *π* for this species. For Greenland halibut (*Reinhardtius hippoglossoides*) and European plaice (*Pleuronectes platessa*) we estimated *π* from publicly available RAD-seq- and whole genome shotgun sequencing data, respectively. The across-species comparison of *π* distribution in MRRs and FRRs revealed a striking agreement among the four species analyzed, with regions corresponding to Atlantic halibut MRR having higher *π* (Fig. 4a-c), indicating that heterochiasmy has been conserved among righteye flounders.

**Fig. 4:**
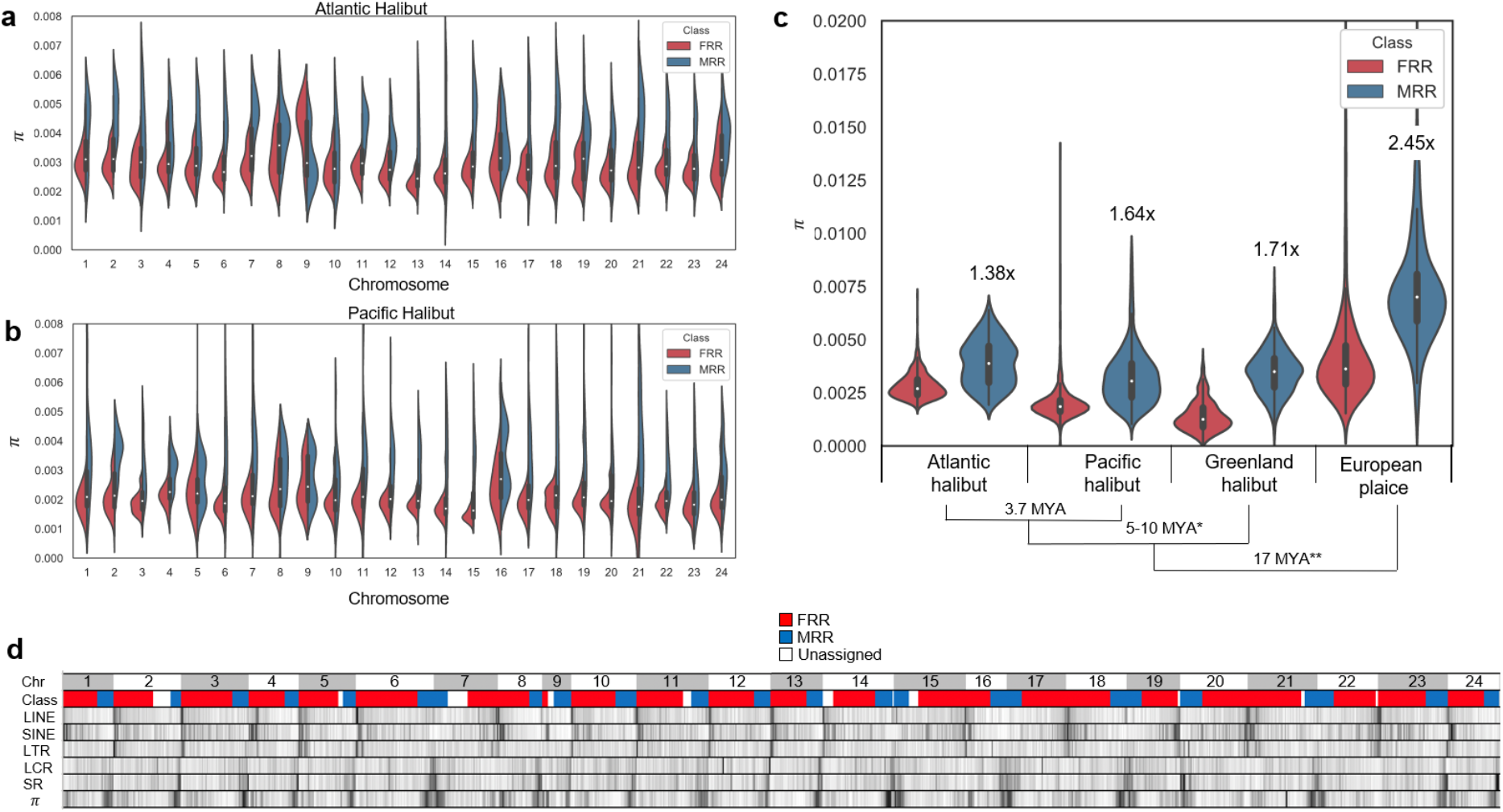
Nucleotide diversity (*π*) and repeat content is correlated with sex-restricted recombination. **a** *π* distribution in the Atlantic halibut genome, contrasting regions of male- vs. female restricted recombination, MRRs and FRRs, respectively. **b** Distribution of Pacific Halibut *π* estimates for regions mapped back to the Atlantic halibut genome, classified as either MRRs or FRRs. Chromosomal and MRR/FRR assignments were extracted from the Atlantic halibut genome assembly coordinates. **c** Genome-wide distributions of estimated *π* for Atlantic-, Pacific- and Greenland halibut and European plaice in regions corresponding to herein defined Atlantic halibut MRRs and FRRs. Numbers above MRR distributions indicate, within species, how much higher *π* is in MRR regions compared to FRR regions. * The time since divergence of Greenland halibut from *Hippoglossus* was estimated from^24^. ** Divergence estimate for European plaice was obtained from TimeTree^25^. **d** Densities of repeat classes and *π* for 500 kb windows along each Atlantic halibut chromosome. Class color indicates whether a region was annotated as FRR, MRR or was unassigned. LINE and SINE represent Long- and short interspersed nucleotide elements, respectively. LTR indicates Long Terminal Repeats. LCR and SR indicate low complexity- and simple repeats, respectively. *π* indicates nucleotide diversity. The white to grey scale in heatmaps is proportional to the min. and max. values within each class.

We investigated whether MRRs and FRRs differed for repeat content and annotated repeats in IMR_Hiphip.v1 by means of RepeatMasker^26^ using the *Actinopterygii* database. RepeatMasker raw output was parsed by repeat subclass and the density per subclass was investigated for each chromosome (Supplementary Table 4). Overall, Long- and Short Interspersed Nucleotide Element (LINE and SINE) densities increase towards the FRR end, while simple repeats and low complexity repeats show the opposite pattern, being enriched towards the MRR end (Fig. 4d). Chr13 and chr10 (Supplementary Fig. 9 and 10) are both typical with respect to clustering of major repeat classes into MRRs/FRRs. Chr10 is the only example where the MRR end overlaps the predicted centromere, suggesting that the clustering and accumulation of repeats is shaped by evolutionary forces associated with sex restricted recombination rather than centromere location.

### The insertion on chromosome Y is of transposon origin and contains a *gsdf* promoter

In an effort to identify candidate Structural Variants (SVs) between Y and X that may influence *gsdf* expression, we aligned all the ONT reads from the male individual used for genome assembly to the IMR_hipHip.v1 assembly and used the software Sniffles^27^ to call SVs along the genome. This analysis revealed a large set of putative SVs (Supplementary Table 5), two of which coincided with the *gsdf* locus, both detected as heterozygous deletions in relation to the reference at chr13:8503901-8505128 (1,227 bp) and chr13:8509688-8509767 (79 bp). Manual inspection of these hits revealed that both ONT reads and linked read pseudo-haplotypes from the male assembly individual validate the assignments made by Sniffles. We used alignments of the male and female pool sequencing data and the phases of ONT long reads to conclude that the insertion allele at these two indels is present on chrY. The HiC data used for scaffolding the assembly was used to screen for large-scale SVs that may differ between X and Y. This analysis indicated that no large inversions are present on chr13 and that neither Y nor X are part of any large translocations with other chromosomes, as such events would appear as off-diagonal contacts in the contact map (Supplementary Fig. 11).

The 1.2 kb insertion upstream of *gsdf* (INS1) is not found in the corresponding Pacific halibut locus, supporting that this region is derived on the Y chromosome in Atlantic halibut. INS1 (IMR_Hiphip.v1 chr13:8503902-8505127) is highly similar to 24 other loci in the genome assembly. Sequence alignments showed high sequence similarity starting with TG and ending with CA di-nucleotides in 19/24 loci. The remaining 5 loci showed strong, further extending similarities to each other, and we therefore extracted additional sequence flanking these five loci to perform a 7-8 kb alignment, which also terminated in conserved TG/CA di-nucleotides in four out of the five sequences. The TG/CA_-e_nds are hallmarks of retrotransposon long terminal repeat (LTR) boundaries, and we could also identify 4-6 nucleotide sequence duplications immediately flanking the insertions, indicating target site duplications (TSDs) that confirm the transposable element (TE) insertions as retrotransposons. Although these TE hallmarks were obscured at the INS1 site upstream if *gsdf*, the sequence similarity and alignment to the related loci in the genome support its TE origin (Supplementary File 1). One of the five longer sequences (IMR_Hiphip.v1 chr14:3028070-3045206) was used as query to search the DFAM database^28^. This search revealed a 5254 nt long hit against the 5410 nt long *Danio rerio* Gypsy transposable element (Gypsy6-I_DR, Bit score 2342), supporting by sequence similarity association that INS1 was derived from a Gypsy retrotransposon that inserted in antisense orientation relative to *gsdf*. Strong similarities between the full-length Gypsy LTR sequences and INS1 revealed that INS1 is indeed an LTR remnant of an ancient Gypsy insertion. This association is further supported by RepeatMasker analysis of the five putative full-length hits partially matching INS1, as they all contained Gypsy sequence annotations.

From the ONT data used for genome assembly we could assess DNA methylation at the *gsdf* locus. Interestingly, we observed low DNA methylation within INS1, in a region coinciding with a predicted CpG-island (Fig. 5 and Supplementary Fig. 12). Further inspection of the lowly methylated CpG-island revealed predicted Sp1 binding sites indicating that this locus may act as a promoter region for *gsdf*. It is clear from both ONT read alignment and the linked read pseudo-haplotype alignments that the chrY haplotype indeed carries INS1. We developed a PCR assay to amplify into INS1 and observed that this assay gave clear bands of the expected size for 10 adult phenotypic males while no bands were observed for 10 phenotypic females (Supplementary Fig. 13). Expression of embryonal *gsdf* (28-54 dpf) from the Y-allele (*gsdf*-Y) extends further upstream of the transcription start site than gene models both in adult Atlantic halibut testis and in male- and female reproductive tissues in adult Pacific halibut. The longer *gsdf-Y* isoforms contain 5’UTRs extending into the Gypsy LTR INS1 (Fig. 5 and Supplementary Fig. 12), which supports promoter activity of INS1.

**Fig. 5:**
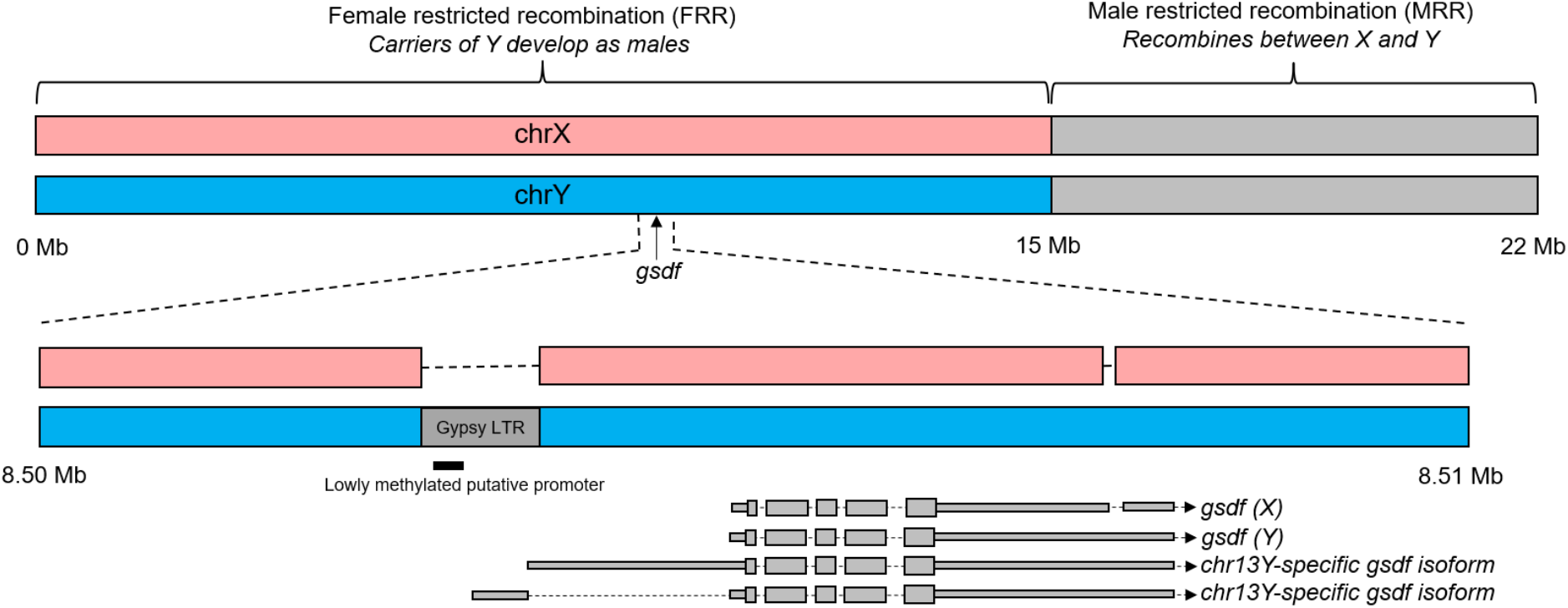
Chr. Y carries a transposon-derived LTR upstream of *gsdf*. Schematic representation of findings. Male and female chr13 haplotypes differ for a 1.2 kb transposable element derived segment 2 kb upstream of the *gsdf* transcription start site. Males carry the insertion allele which contains a lowly methylated CpG-island inferred to act as a derived promoter of *gsdf* uniquely in males. The shorter *gsdf* isoforms show the extent of *gsdf* expression observed in adult testis in Atlantic halibut as well as adult ovary and testis of Pacific halibut. The two *gsdf* gene models labeled chr13Y-specific indicate isoforms detected only in males from three separate developmental stages. An IGV-plot of this region is presented in Supplementary Fig. 12.

### Analyzing chrY decay

In addition to the ONT genome assembly, we also sequenced the same male using a Chromium Genome linked read library and used the resulting data to generate two separate pseudo-haplotype assemblies which successfully phased the first 10 Mb of chr13 into separate scaffolds for the X and the Y chromosome. Loss of X/Y recombination is expected to lead to chrY decay, and we investigated whether we could find evidence of this using the phased chrX (X^PH^) and chrY (Y^PH^) along the first 10 Mb of chr13. This analysis, which involved quantification of possibly deleterious variants along the first 10 Mb of X^PH^ and Y^PH^, revealed a 30% excess of variants predicted to be deleterious (HIGH classification in SNPEff) on Y^PH^ (n=35) compared to X^PH^ (n=27). That no large-scale loss of genetic material has occurred on chrY is supported by the observation that super-scaffolds of X^PH^ and Y^PH^ were both 20.4 Mb in size.

## Discussion

In this study we show that *gsdf* is most likely the master sex determining gene. Our interpretation is that following an ancient Gypsy element insertion on the autosomal progenitor of chr13, this element degraded and left a remnant LTR, which now acts as a promoter driving the expression of *gsdf* ectopically during development of XY individuals, making carriers develop into phenotypic males. Interestingly, In Luzon ricefish, where *gsdf* is also the MSD gene, the region immediately upstream of *gsdf* was shown to be most important for modulating *gsdf* expression^22^. The region tested corresponds approximately to the site of TE insertion in Atlantic halibut. A role of *gsdf* in early gonadal development is supported by experiments in medaka where targeted homozygous deletion of *gsdf* in genetic (XY) males triggers an early development of ovaries, with the majority of these *gsdf* KO homozygote fish still developing testes later in life^29^. It would be interesting to use CRISPR/Cas9 to remove the Gypsy LTR from XY Atlantic halibut (X/Y^LTR-^) and insert the Gypsy LTR into XX individuals X/X^LTR+^), as we have done in Atlantic salmon^30,31^. Our predictions are that X/Y^LTR-^ individuals would develop as females and X/X^LTR+^ individuals as males. Furthermore, in accordance with previous studies^32^, we observe *gsdf* expression in adult testes and ovaries in Pacific halibut and in Atlantic halibut testis, indicating a function of Gsdf in mature gonads. In medaka ovaries and testes, *gsdf* is expressed by somatic cells surrounding germ cells, with higher expression in testis^29^. In Atlantic salmon *gsdf* expression has been detected in Granulosa and Sertoli in ovaries and testes, respectively^33^. This supports a role for Gsdf in the mature gonad and suggests that high expression of *gsdf* from Atlantic halibut chrY during development is expected to trigger testis development. TEs are known to be involved in sex-chromosome evolution^34^, e.g. TEs are probably responsible for translocation of *sdY* in salmonides^35^ and in Medaka a LINE insertion at the *dmrt1b* locus has been implicated as the sex-determining variant^36^. It has been repeatedly observed that TEs are repurposed for genomic innovation by domestication^37^, as might be expected for a dynamic part of the genome.

In this study we observed that meiotic recombination is dramatically different between male and female gonads, and this can also be seen when revisiting a previous Atlantic halibut linkage study^20^, where this phenomenon was not strongly emphasized. Interestingly, the guppy (*Poecilia reticulata*) has genome-wide heterochiasmy, while no significant difference was observed in the closely related genus *Xiphophorus*^38^, suggesting that the mechanism underlying heterochiasmy may be easily modified over short evolutionary time-spans. A common feature for observations of heterochiasmy in teleosts is enrichment of male crossovers towards chromosome ends^38-40^, which is also the case in Atlantic halibut. We found that four analyzed righteye flounder species all had higher *π* in the regions orthologous to Atlantic halibut MRRs than in corresponding FRRs orthologous regions (Fig. 4c). This likely reflects conservation of sex restricted recombination across *Pleuronectidae*. The reason for this interpretation is that few polymorphic sites are expected to be shared between species over the millions of years separating the four analyzed species. Thus, the diverging *π* distributions must have been established independently, presumably by selective forces associated with heterochiasmy.

In medaka, sex restricted recombination between sex chromosomes has been shown to follow the XY male pattern also in XX neomales^41^, which supports that a mechanism associated with the gonadal environment governs recombination suppression in meiosis. We propose to study the mechanisms of heterochiasmy in sex-reversed genetic female Atlantic halibut, which are used as breeding males in aquaculture. Regions of TE-accumulation in genomes have been implicated as putative heterochromatin sinks^42^, and both TEs and heterochromatin are positively correlated with suppressed recombination^43^. It is difficult to disentangle whether TEs are involved in the establishment of meiotic recombination suppression or if their differing distributions in MRRs/FRRs is merely a consequence of how the genome evolves due to unknown selective pressures imposed by heterochiasmy already being in place. One possibility could be that Atlantic halibut male and female gonads differ for which regions of the genome become heterochromatin, perhaps as a defense against TE transposition, and that male and female gonads direct recombination differently in response to variable TE content along the chromosomes. Possibly, transposable elements are more likely to accumulate in FRRs, for example if regions only recombining in males (MRRs) exist in a state less accessible for transposition during female and/or male meiotic recombination. Accumulation of local TE clusters in FRRs may also contribute to the further homogenization of this chromosome compartment by gene conversion of homologous TE insertions. It should be noted that we have analyzed repeats broadly, and future studies aiming to investigate repeat classifications in relation to heterochiasmy may benefit from more detailed classifications. We show that *π* is higher in MRRs than in FRRs and hypothesize that the higher recombination rate on the physically smaller MRRs increase the probability of retaining diversity at linked sites during purifying selection^44^, or that differences in purifying selection due to the differing TE-composition landscapes of FRR and MRRs has been involved in shaping *π*.

Sex-restricted meiotic recombination, as we describe here, suppresses recombination for large segments along autosomes. Due to genome-wide, pre-existing sex-restricted recombination being in place, this system is expected to facilitate the survival of novel sex determining factors which in turn could give rise to new, and even competing genetic sex determination systems (Supplementary Fig. 14). We propose that heterochiasmy has the potential to facilitate evolution of sex determination systems, and presumably speciation as postulated in stickleback^45^. It has been reported that female is the heterogametic sex (ZZ/ZW) in both Pacific halibut and European plaice^12^ as well as the half-smooth tongue sole^46^, with the main candidate MSD gene being *dmrt1*. This makes it conceivable, but not certain, that the Atlantic halibut X/Y system replaced a Z/W system shared across different flatfish species.

Regardless of heterochiasmy being a facilitating factor in the evolution of sex determination systems or not, we conclude that the TE insertion in the *gsdf* locus occurred after the split between Atlantic halibut and Pacific halibut around 3.7 MYA. Sex chromosomes originate from autosomes that acquire a sex determination locus and then lose recombination with the former autosomal pair, which typically involves structural variation in the non-recombining region of the sex chromosome pair, while other regions may still recombine and act as pseudo-autosomal regions. For mammalian Y, as well as avian W, loss of purifying selection over millions of years has led to pseudogenization, gradual loss of genes and a reduction in chromosome size^47^, where remaining genes and regulatory elements determine male and female phenotypes, respectively. Our analysis of deleterious alleles identified a non-significant 30% increase in predicted deleterious alleles on chrY compared to chrX, which may indicate an early stage of chrY decay. We observe a 1 Mb segment (chr13:5.5-6.5 Mb) (Fig. 1c) showing differentiation equal to the genome average, interpreted as X/Y recombination, although rare, not being fully suppressed in the FRR of chr13. This is supported by previous fine mapping of the sex determining locus to a smaller marker interval^21^ (Fig. 3a) in a region corresponding to the herein defined chrY. A retrotransposon insertion into the *gsdf* locus started the transformation of an autosome into a sex chromosome in Atlantic halibut less than 4 million years ago. Incomplete recombination barriers and little, if any, chrY decay both suggest that Atlantic halibut chrY is in the early stages of the evolutionary trajectory of a sex-chromosome.

## Methods

### Ethics statement

For this study the halibut were reared under conditions similar to standard commercial fish farming conditions. Such conditions are listed as an exception in The Norwegian Regulation on Animal Experimentation, thus approval of the experimental protocol of this experiment by NARA (the governmental Norwegian Animal Research Authority) was not required.

### Halibut material

All samples were taken during routine production of Atlantic halibut at the Austevoll Research Station, Institute of Marine Research, Storebø, Norway. Halibut breeding individuals were stripped, eggs fertilized and incubated, and juvenile halibut produced according to standard aquaculture protocols developed at the Austevoll Research Station^48-50^. Briefly, females were stripped according to their individual ovulatory rhythms to ensure high (>90%) fertilization. Fertilized eggs were incubated in 250 l conical incubators with upwelling water for 60 day degrees (i.e. 10 days at 6°C) and transferred to larger incubation units, silos (vol. 50,000 l) before hatching. Yolk sac larvae were held in the silos until ca. 265 day degrees posthatch when they were moved to first feeding tanks and fed *Artemia* for 70 days until metamorphosis was complete. At this stage, the larvae were weaned onto a formulated diet. Juvenile halibut were moved to ongrowing tanks when they had reached an average size of ca. 10 cm and held there until they had reached a harvest size of 2-5 kg. One adult, three-year old male was euthanized with metacain (100 mg/l) and muscle, spleen and blood samples were taken for reference genome sequencing. For sequencing of pooled family samples, tissue samples were taken from ten full siblings of mixed sex (ten males and ten females). All samples were snap frozen in liquid nitrogen (N_2_) and stored at -80°C before DNA extraction. RNA-sequencing was made on a series of samples taken through embryonic development. Before hatching, embryo samples were snap frozen in liquid N_2_ and stored at -80°C. Larvae were euthanized in metacain (100 mg/l), transferred into RNA later, kept at 4°C for 24 h and stored at -20°C until RNA isolation. The samples for qPCR were snap frozen in liquid N_2_ and stored at -80°C. Digital images of all embryo-larval stages were captured using video cameras Moticam 1080 (Motic) mounted on Olympus SZX10 stereomicroscope using Motic Live Imaging Module. Living embryos and larvae were individually immobilized in a Petri dish containing 2% methylcellylose/98% seawater. Images were then scaled and processed by ImageJ software^51^.

### DNA extraction and sequencing of male and female pools

DNA was extracted from muscle samples using Qiagen DNeasy Blood and Tissue Kit. Paired-end libraries of equimolar male (10 individuals) and female (10 individuals) DNA pools and from one single individual of each sex were constructed using the Genomic DNA Sample Preparation Kit (Illumina) according to manufacturer’s instructions and sequenced on the Illumina HiSeq2000 platform (Illumina) at the Norwegian Sequencing center (https://www.sequencing.uio.no, Oslo, Norway).

### High Molecular Weight DNA extraction and sequencing

Several tissues were collected from a single male Atlantic Halibut. Muscle and spleen were snap frozen in liquid nitrogen and stored at -80°C until further use. Fresh blood was used to isolate erythrocyte nuclei and then prepare High Molecular Weight (HMW) DNA from these nuclei according to a Phenol/Chloroform protocol (Quick). Extracted DNA was either left unfragmented or sheared using either MegaRuptor (Diagenode) or by passing DNA through a 100µl pipette tip 10-20 times. From these DNA samples we produced MinION sequencing libraries with the standard adapter ligation protocol kit (LSK-108), 1D^2 sequencing kit (LSK-308) and the RAD002 and RAD003 kits (Oxford Nanopore Technologies). For libraries prepared with the RAD002 and RAD003 kits we used a protocol developed to produce ultra– long reads (Quick, protocols.io). In total we used 9 flow cells and sequenced 18.8 Gbp in 2,963,108 reads with a read N50 of 17.8 kb. Sequenced fast5 raw signal data files were basecalled using albacore v.2. The size distribution of sequenced reads is presented in **Supplementary Figure 1a**. We also used the collected muscle samples to produce HiC libraries (Dovetail), sequenced on one lane of HiSeq X (Illumina). One of the DNA preps inferred to have HMW DNA was used to prepare a linked reads Chromium library (10x genomics) and was sequenced on one lane of HiSeqX (Illumina).

### Genome assembly, polishing and scaffolding

Nanopore reads passing filters were used in assembly by first constructing all-vs-all alignments between all reads using minimap2^52^. These all-vs-all alignments were then used in the software miniasm to produce a haploid contig assembly. The contig assembly was polished using racon^53^ with the nanopore data followed by pilon^54^ using the Illumina data from the male pool. Scaffolding of the polished miniasm assembly was performed using the HiRise pipeline (Dovetail Genomics). The quality of the generated assembly was assessed using BUSCO^55^. Initial nanopore contigs assembled with miniasm^52^ had a contig N50 of 3.2Mb and the resulting final assembly had a scaffold N50 of 27.2 Mb.

### Assembly of linked read data

The software Supernova2^56^ (10X Genomics) was used to produce a genome assembly of the sequencing data resulting from the Illumina sequencing of the Chromium library. For this assembly, reads were subsampled to get close to the recommended 56x genome coverage recommended in the Supernova2 user manual. The two pseudo-haplotype linked read assemblies featured one scaffold each which covered the whole of IMR_Hiphip.v1 chr13. We used SNP markers identified as divergent between the male and female short read data to investigate the pseudo-haplotypes in relation to sex divergent markers, i.e. if one pseudo-haplotype could represent the Y chromosome and the other the X-chromosome. This analysis revealed that for the first 10 Mb of chr13, the allele being heterozygous in males and not present in females (the perceived Y-allele) was observed on pseudo-haplotype number 2 (1,954 SNPs). The perceived Y-allele was observed only for 2 SNPs in this region for pseudo-haplotype number 1. The corresponding counts for the rest of chr13 was 125 and 213 alleles on pseudo-haplotype numbers 2 and 1, respectively.

### RNA-sequencing and gene annotation

Total RNA was extracted from whole larvae from seven different stages of development using miRNeasy Mini Kit (Qiagen) and treated with TURBO DNase-Free kit (Ambion). TruSeq cDNA libraries were sequenced on the HiSeq 4000 sequencing platform (Illumina). The average number of reads was 50,320,639 per sample. Twenty-two samples (four samples from 54 dpf and three samples from each other stage) were assembled individually with Trinity using the Agalma pipeline^57^. The transcriptome assemblies were annotated with BLAST against Swissprot (download date 20191227) and the Ensembl gene models of zebrafish (GRCz11) and stickleback (BROAD S1). The hits with the best scores were kept. Transcripts from all assemblies were mapped to the halibut genome with BLAT. The thresholds for mapping were set to 98% match for sequences with length >=401 nt and 95% match for sequences with length <=400 nt. Long sequences were mapped before short sequences. The mapped sequences were clustered on genome location, selecting the sequence with the highest score for each non-overlapping cluster. Only clusters with an annotation (>300 in BLAST score threshold) were saved. Neighboring clusters annotated with the same gene from either Swissprot, zebrafish or stickleback were merged in cases where the annotation indicated that the gene was split in several clusters (the neighboring clusters were only merged in cases where the annotation was from different parts of the same gene). For the merged clusters, the sequence with the highest BLAT score from all merged clusters were chosen. Additionally, the transcriptome assembly sequence with the highest scoring BLAT match to the merged cluster was added. We found 15,273 annotated genes based on the Trinity assemblies. The zebrafish and stickleback ENSEMBL gene models were mapped to the halibut genome using exonerate in order to add genes without RNA-seq evidence. The exonerate hits were clustered by location, using the exonerate hit with the highest score for each cluster. Sequences for each exonerate based gene were extracted from the halibut genome sequence. 2,269 genes were added based on the exonerate analysis. We added 222 genes that had no annotation (BLAST threshold score <300) but had RNA-seq expression. After running BUSCO 3.1 with the Actinopterygii odb9 dataset on the non-filtered Trinity assemblies that mapped to the genome^55^, we added 550 genes from the BUSCO analysis that were missing from our filtered list. This resulted in a total number of 18,314 genes and 20,579 transcripts giving a 80.9% complete, 8.7% fragmented and 10.4% missing BUSCO coverage.

### Nanopore sequencing of testis RNA

Testis tissue was homogenized at 4°C in 1ml QIAzol Lysis Reagent (Qiagen, Hilden, Germany) together with RNase-Free Zirconium Oxide Beads (NextAdvance, Inc., Troy, NY, USA) during 1 min, maximum effect, in a BulletBlender (NextAdvance) and total RNA was prepared according to the QIAzol (Qiagen) protocol. Polyadenylated RNA was isolated with the Dynabeads mRNA direct purification kit (Thermo Fisher Scientific), following the kit protocol. For full-length cDNA Oxford Nanopore sequencing, the cDNA-PCR Sequencing SQK-PCS108 kit (Oxford Nanopore Technologies) was used according to the kit protocol with minor modifications. The finished cDNA library was sequenced on a MinION device using a R9.4 flow cell (Oxford Nanopore Technologies).

### Screening for male vs. female divergence

Illumina sequences from the male- and female DNA pools were filtered by trimmomatic^58^, then aligned to the assembly IMR_hipHip.v1 using BWA-MEM^59^. Aligned reads were subjected to duplicate flagging (“Picard Toolkit.” 2019. Broad Institute, GitHub Repository. http://broadinstitute.github.io/picard/; Broad Institute) and were then used as input for calling SNPs as well as small Insertions/Deletions using Genome Analysis Toolkit (GATK)^60^ UnifiedGenotyper. Raw variant vcf files were subjected to variant filtration using Best Practices procedure with SNP-clusters being added as an additional filter (more than 5 SNPs in a window of 20 bases were filtered out). SNPs passing filters were isolated from the vcf file and allele frequencies from the male- and female pools were extracted and used in *χ*^2^-analysis examining the null hypothesis; that allele frequencies were not different between the male- and female pools. Prior to the *χ*^2^-test, in order not to inflate the significance at positions with high depths of coverage, SNP read depths >12 for either pool were set to 12 while retaining the observed allele frequency ratio. *χ*^2^-tests were then performed SNP-by-SNP, and average *χ*^2^-scores of non-overlapping windows of 100 adjacent SNPs along the chromosomes were extracted.

### Sex differentiating PCR assay

Genetic sex of each embryo was determined by PCR using sex-specific primers (Table 1). The thermal cycling conditions were as follows: 94°C for 5 min, 32 cycles with 94°C for 30 sec, 65°C for 1 min and 72°C for 1 min, and finally 72°C for 5 min. Each 12 µl PCR reaction contained 5X GOTaq Flexi Buffer, MgCl, dNTPs, GoTaq Polymerase, PCR primers and template cDNA.

**Table 1.**
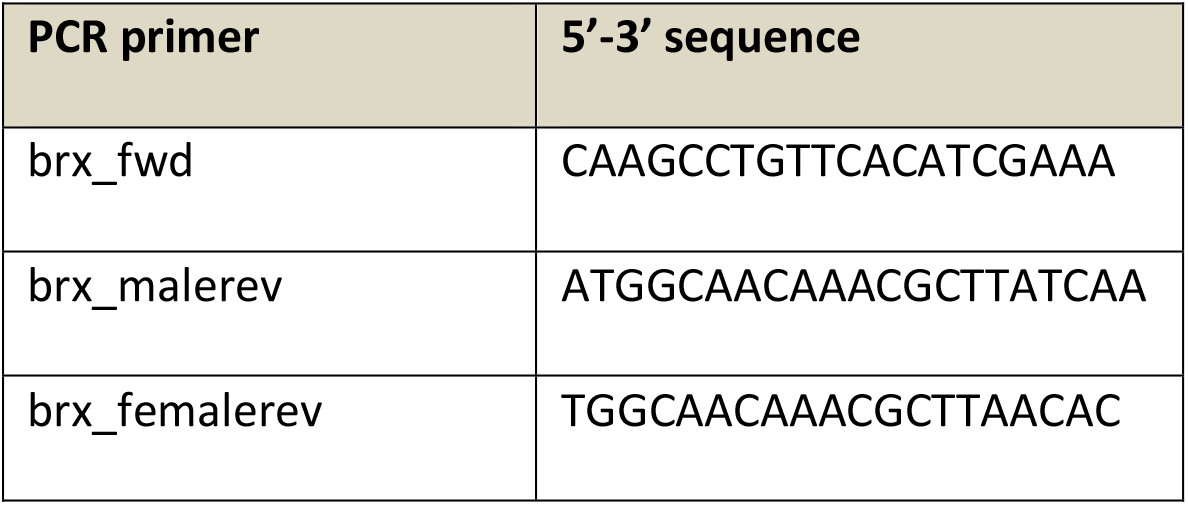
Brx PCR primers.

### Differential gene expression between sexes

We generated RNA-sequencing data from 12 Atlantic halibut whole embryos representing stages 28 (n=4), 43 (n=4) and 54 (n=4) dpf. Since one cannot determine the phenotypic sex at this stage we aligned the RNA-seq data using Bowtie 2^61^ to IMR_hipHip.v1 and extracted expressed allele counts at positions identified as divergent between males and females in the pool DNA-seq. Extracted RNA-seq allele ratios were used to cluster the individuals, revealing two well-separated clusters of XY and XX individuals and this assignment was used to identify genetic male and female individuals for the differential expression screen. The differential expression screen involved contrasting across-sample normalized gene expression levels of the seven inferred males to the five inferred females in a two-tailed t-test. Principal Component Analysis (PCA) was performed, using the R function “prcomp”, on the normalized gene expression levels from the 12 embryos.

### *gsdf* quantitative PCR

Genomic DNA and total RNA was extracted from single embryos using the Allprep DNA/RNA/miRNA Universal Kit (Qiagen), according to the manufacturer’s instructions. The RNA was DNase treated as part of the extraction process, according to the manufacturer’s instructions. For each sample, cDNA was synthesized from 125 ng RNA using the Superscript VILO cDNA synthesis Kit (ThermoFisher Scientific). The primers used for qPCR were designed using BatchPrimer3 (https://wheat.pw.usda.gov/demos/BatchPrimer3/) (Table 2). The gene *gtf3c6* was used as an internal reference gene for normalization of the qPCR data due to its stable expression in the developmental stages analysed in this study (Supplementary Fig. 15). Using different RNA concentrations, the PCR efficiencies (E = -1+10(−1/slope)) for the primers were calculated to be 108.5% (*gsdf*) and 92.4% (*gtf3c6*). A melt curve analysis was performed to confirm that the primers did not amplify more than one PCR product. qPCR was performed in duplicates in 384-well optical plates in a QuantStudio 5 Real-Time PCR system (ThermoFisher Scientific) using default settings. 2 µl cDNA (diluted 1:20) was used in a 6 µl Fast SYBR Green qPCR reaction (ThermoFisher Scientific). No-template controls for each gene were run in all qPCR plates. The relative gene expression level was calculated using the comparative Ct (or 2^−ΔΔCt^) method. All values were normalized to *gtf3c6* and calibrated to the sample with the lowest ΔCt.

**Table 2.**
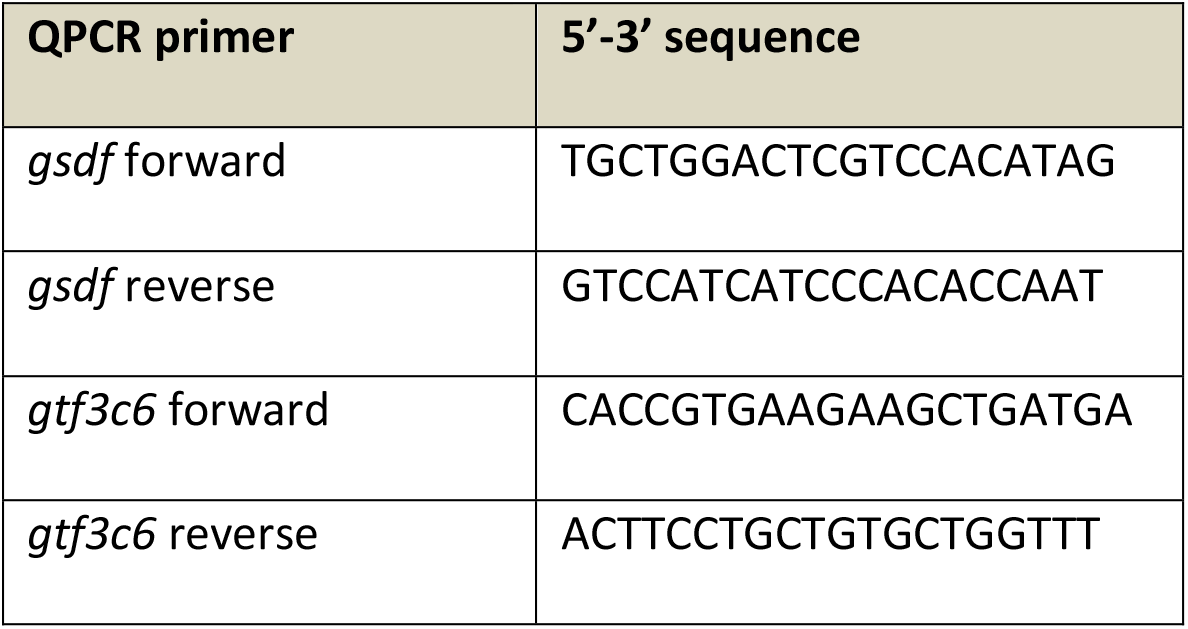
Primers used for qPCR.

### Detection of structural variants, small insertion/deletions and annotation of repeat sequences

The long-read SV identification tool Sniffles^27^ was used to identify SVs from the ONT data. The ONT data from the single male individual used for genome assembly was aligned to IMR_hipHip.v1 using minimap2^52^ with the “-x map-ont” option and the resulting bam file was used in sniffles, requiring >=5 observations of a variant allele to be retained as a potential SV.

### Analysis of chrY decay

The two linked-read pseudo-haplotype assemblies of the male Atlantic halibut used for the genome assembly were aligned against the Pacific halibut reference genome (nucmer) and SNPs and small indels were called using the “show-snps” function of the MUMmer 4.0 package. Identified variants were converted to vcf-format (https://github.com/MatteoSchiavinato/Utilities/blob/master/my-mummer-2-vcf.py) and were then annotated using SNPEff, with gene models derived from the Pacific halibut IPHC_HiSten_1.0 gtf-file “GCF_013339905.1_IPHC_HiSten_1.0_genomic.gtf”. SNPs and indels that were unique to either pseudo-haplotype were investigated for their annotations. Annotation frequencies were compared between the two pseudohaplotypes for the interval where they represent the X and Y (Atlantic halibut: chr13:0-10Mb, Pacific halibut: NC_048944.1:0-11Mb).

### Analysis of the evolutionary origin of the 1.2 kb insertion upstream of *gsdf*

The 1.2 kb insertion (IMR_Hiphip.v1 chr13:8503902-8505127) was used as a query sequence in BLAT^62^ searches against IMR_HipHip.v1. Full length hits were retained (n=30) and were extended to include 2kb up- and downstream sequences and these approximately 5.3 kb sequences were used as input for multiple sequence alignment using MUSCLE^63^. The resulting alignments were visualized in AliView^64^ and abrupt drops in sequence similarities were observed for 24 of the sequences on each flank. The extent of highly similar sequences started and ended at perfectly conserved TG and CA di-nucleotides, the hallmark motifs of LTR boundaries, and the insertion was immediately flanked by 4-6 nucleotide duplications of loci TE target sites (TSD) confirming the LTR-retrotransposon insertion.

### DNA methylation calling from ONT data and CpG island prediction

Basecalled ONT reads previously used for the genome assembly were aligned back to the IMR_Hiphip.v1 assembly using minimap2. The software nanopolish^65^ was used to index the raw ONT fast5-files to enable extraction of likelihoods of CpG site methylation along individual mapped reads in genome assembly coordinates. Next, the nanopolish script “calculate_methylation_frequency.py” was used to extract the methylation frequency at each adequately covered CpG cluster in the assembly. CpG islands were predicted for the IMR_Hiphip.v1 assembly using gcluster^66^ employing default settings.

### Estimation of nucleotide diversity

For Atlantic Halibut we estimated *π* by creating chromosome-specific samtools mpileup files from the aligned data previously used for variant calling, using only positions with a mapping quality >=20 and read depths ranging between 10 and 50. The script “WinNuclDivmPileup.py” (https://github.com/cjrubinlab/python_scripts2) was used to estimate *π* in 500kb windows along the genome for the male- and female pools. For Pacific halibut no DNA-sequencing data was available in public sequence repositories, so we devised a way to estimate *π* from RNA-sequencing data. Sequencing data from 12 RNA-seq experiments of various tissues and individuals was downloaded from the SRA (SRR11826178-SRR11826189) and these data were aligned to the Oxford nanopore derived Pacific Halibut genome assembly (GCF_013339905.1) using the STAR aligner^67^. The resulting alignments were randomly subsampled 20 times and then subjected to duplicate-pair removal using picard-tools MarkDuplicates, retaining 70% of the reads after filtering. From the bam file containing filtered reads an mpileup file was generated using samtools (mapping quality exclusion filter < 20). For the Greenland Halibut (*Reinhandtius Hippoglossoides*) we downloaded the genome assembly from NCBI (GCA_006182925.2_MU_Rhippo_v1) as well as RAD-sequencing data from 20 individuals (SRA accessions: SRR11351808-SRR11351827), aligned the short-read data to the assembly using minimap2 with the “-x sr” option. From mpileup files derived from samtools mpileup for Pacific Halibut and Greenland halibut the estimated heterozygosity (He) was calculated for positions having coverage>=10 and mapping quality >=20 using the WinNuclDivmPileup.py” script. For European plaice (*Pleuronectes platessa*), a species lacking a genome assembly, we downloaded paired genome resequencing Illumina reads (SRA accession: SRX8240416), aligned these reads to the IMR_Hiphip.v1 assembly using bwa and proceeded with the same pipeline as was used for estimation of Atlantic halibut *π*. In order to lift over genomic intervals annotated as MRRs and FRRs in the Atlantic halibut to the Pacific- and Greenland halibut assemblies we used the satsuma^68^ commands “Chromosemble” to generate a detailed list of lift-over intervals and “BlockDisplaySatsuma” to summarize the detailed list into coherent blocks. We then converted these coordinates to the bed-format and used the bedtools^69^ “intersectbed” command to link coordinates in the other assemblies to the corresponding coordinates in the Atlantic halibut assembly and MRRs and FRR classification.

### Linkage analysis

RAD-sequencing data from an Atlantic halibut half-sib pedigree altogether comprising three parents and 90 offspring were downloaded from the SRA (accession=SRX193134). The downloaded individual files were split by barcodes and were then connected with annotation information from the original publication regarding phenotypic sex, parent/offspring status and family B/C^21^. The paired-end data from each individual in the pedigree was then aligned to IMR_hipHip.v1 using BWA-MEM. Resulting bam-files were subjected to duplicate removal using picard-tools “MarkDuplicates”, then raw variant (SNP and indel) calling using GATK UnifiedGenotyper. Raw SNPs were filtered using GATK “VariantFiltration” with the following arguments, also with a list of raw indel positions given to the “InDel” mask; “--maskName InDel”, “--maskExtension=5”, “--clusterSize=5”, “--clusterWindowSize=15”, “BaseQRankSum<-7.0”, “BaseQRankSum > 7.0”, “MQRankSum < -7.0”, “MQRankSum > 7.0”, “QD < 5.0”,“FS > 50.0”,“ReadPosRankSum > 7.0”, “ReadPosRankSum < -7.0” and “DP > 40”. SNPs remaining after filtration were saved in a new vcf file which was given as input to Lep-MAP3^70^ to generate recombination maps for the two families. The commands used in Lep-MAP were: “ParentCall2”, with the options (removeNonInformative=1 XLimit=2 halfSibs=1), then “Filtering2”, with the option data=- (dataTolerance=0.01), then “SeparateChromosomes2” with the options (lodLimit=10 distortionLod=1) to identify linkage groups in the pedigrees. SeparateChromosomes2 with lodLimit=10 generated a map of 31 linkage groups, out of which 20 had 964–3063 markers assigned to them (24 were expected based on karyotype). We continued with the output from SeparateChromosomes2 (lodLimit=10) to run “OrderMarkers2” to, for each identified linkage group, order the markers without having given the software access to genome assembly coordinates. The output files from the first iteration of “OrderMarkers2” were then subjected to a second round of “OrderMarkers2”, this time giving the Lep-MAP3 access to genome coordinates of markers on each linkage group using the option “evaluateOrder”. The resulting linkage group specific files containing male- and female centiMorgan (cM) positions of each marker were filtered to prune obviously inflated cM positions in the very beginning and the very end of assembled chromosomes and by splitting four linkage groups where two assembled chromosomes had been merged into one linkage group, thereby increasing the observed linkage group count from 20 to the expected 24. Resulting maps from this stage were retained for downstream analyses.

### Pacific halibut RNA-seq contrasts

Sequencing data from 12 RNA-seq experiments of various tissues and individuals (SRR11826178-SRR11826189) was downloaded from SRA. Four of these tissues (testis, ovary, white muscle and brain) were used for characterization of *gsdf* gene expression levels and validation of gene models from the gff-file accompanying the Pacific halibut assembly on NCBI. Alignments were performed using STAR as described above for *π* estimation.

### RepeatMasker analysis

RepeatMasker v4.1.0^26^ was used to annotate both simple and interspersed repeats. We deployed the RMBLASTN (v2.6.0.+) search engine to scan for known repeats among ray-finned fishes (“-species actinopterygii”) using a database that combined Dfam_3.1 and RepBase-20181026 repeat sequences. A custom Perl script was used to parse the main output table from RepeatMasker and compute the densities of major repeat classes (i.e. “Simple_repeat”, “LINE”, “LTR”, “Low_complexity”, “DNA”, “RC”, “SINE”, “Satellite”, “Unknown”, “Retroposon”) in windows of 50 kbp and 500 kbp, respectively.

## Supplementary data

Supplementary File 1: Integration site analysis of the 1.2 kb insertion upstream of *gsdf*, which controls male development, and related insertions in the Atlantic halibut genome.

## Acknowledgements

We would like to thank the National Genomics Infrastructure/SNP&SEQ Technology Platform in Uppsala for preparing and sequencing the 10X Genomics Chromium Genome library and thank the Norwegian Sequencing Centre (Oslo) for Illumina sequencing of DNA and RNA. Computations and data handling were enabled by resources provided by the Swedish National Infrastructure for Computing (SNIC) at Uppmax, Uppsala, Sweden, with these resources being partially funded by the Swedish Research Council through grant agreement no. 2018-05973. We would like to thank Leif Andersson for comments on an earlier version of the manuscript. We are grateful for financial support from Sterling White Halibut for the sex determining factor part of this work. This study was co-funded by the Norwegian Ministry of Trade, Industry and Fisheries. This research was supported by grants from the Swedish Research Councils Formas (2012-00740 to C.J.R.; 2018-01008 to P.J.; 2017-00413 to A.W) and Vetenskapsrådet 2018-03017 to P.J.; 2018-04444 to A.W.).

## Author contributions

R.B.E., C.J.R and B.N. designed the project. T.H., R.M.J. and M.M. contributed with rearing of fish. R.M.J., P.P. and B.N. collected the tissue samples. S.K.O. extracted the RNA. L.K. performed the qPCR. E.K.S. and C.J.R performed the structural variant bioinformatic analysis. C.J.R. and O.W. did HMW DNA isolation, nanopore sequencing and genome assembly. E.S. generated RNA-sequencing data. A.W. and C.J.R. performed analysis of RepeatMasker data. C.J.R performed and P.J. contributed to the TE analysis. C.J.R performed variant calling, comparative genomics- and divergence analyses. R.B.E. T.F and O.W. performed annotation. R.B.E. designed, and L.K. and S.M. tested the PCR sexing assay. R.B.E., C.J.R and B.N. analyzed the results and wrote the paper. All authors read and approved the final manuscript.

## Data Availability

The authors declare that the data supporting the findings of this study are available within the article and its Supplementary Information. All datasets will be made publicly available under bioproject PRJNA680301 upon publication.

