## Supplementary File 1 for "Heterochiasmy facilitated the establishment of *gsdf* as a novel sex determining gene in Atlantic halibut"

**Exact TSD motif match**

```
chr12:1119677-1122729
CACTTTATGCTAA[TGTAGC-GTGGT--GGGAAAGC-insert-AGGGAGCGCGCATCTGGCTACA]CTAAGCTAAGCT
chr5:22462115-22465234
CTGCTTAGCTTAG[TGTAGC-GAGGT--GGGAAAGC-insert-GGGGAGCG---ATCTGGCAGCA]TTAGCTTAGCAT
chr7:24033655-24036861
CACGCTTTACCAG[TGTAGC-GAGGT--GGGAAAGC-insert-AGAG-GCGCGCATCTGGCTACA]CCAGGAAGGGCT
chr10:55838-5561696
ACTAGAATGACAG[TGTAGC-GAGGT--GGGAAAGC-insert-AGGGAGCGCGCATCTGGCTACA]ACAGCAAGTACA
chr14:20239439-20242728
TTTACGCCCCGTG[TGTAGC-GAGGT--GGGAAAGC-insert-AGGGAGCGCGCATCTGGCTACA]CGTGCAAGACTC
chr5:4765600-4768907
TCAGGAAAGCATG[TGTAGC-GAGGT--GGGAAAGC-insert-AGGGAGCGCGCATCTGGCTACA]CATGGAACATCC
chr21:154906-158198
ATGGTAGTCTAAG[TGTAGC-GAGGT--GGGAAAGC-insert-AGGGAGCGCGCATCTGGCTACA]TAAGGATTAACC
chr4:3765140950
TTGTAAATGCATG[TGTAGC-GAGGCAGAGGAAAGC-insert-AGGGAGCGCGCATCTGGCTACA]CATGCATGACCT
```

**One base difference in TSD motif**

```
chr7:24022629-24025927
AACAACGCCAGAA[TGTAGC-GAGGT--GGGAAAGC-insert-AG-G-GAGCGCATCTGGCTACA]CCAGGAAGGGCT
chr20:24763276-24766621
AACAACGCCAGAA[TGTAGC-ATGGT--GGGAAAGC-insert-AGGGAGCGCGCATCTGGCTACA]ATAACATTTTAT
chr10:8501255-8504553
CTTCTACTGTGTG[TGTGGCAGAGGG--GGGAAAGC-insert-AGGGAGCGGCGATCTGGCTACA]TCTGGGAATGCC
chr10:8543673-8546987
AGCTTCTACTGTG[TGTAGC-GAGGT--GGGAAAGC-insert-AGGGAGCGCGCATCTGGCTACA]TCTGGGAATGCC
chr4:19881828-19885131
GTTAACATGTGAA[TGTAGC-GAGGT--GGGAAAGC-insert-AGGGAGCGCGCATCTGGCTACA]TGTGTATCAACA
chr23:21890121-21893419
TATGCAAGCTATG[TGTAGC-GAGGT--GGGGAAGC-insert-AGGGAGCG---ATCTGGCTACA]TCTGCAGAAACA
```

**TSD motif >= 2 base difference**

```
chr13:8502902-8506130 (candidate sex determining mutation)
TTAATATTGCATC[TGTAGC-GAGGT--GGGAAAGC-insert-AGGGAGCGCGCATCTGGCTACA]TCAAATGCATAT
chr14:180799-184097
ACAGACATGCGAA[TGTAGC-GAGGT--GGGAAAGC-insert-AGGGAGCACGCATCTGGCTACA]CAATGTTAACAA
chr14:3028310-3031425
AACAACGCCAGAA[TGTGGC-GAGGT--GGGAAAGC-insert-AGGGAGCGCGCATCTGGCTACA]CACACATTCAGC
chr20:27177333-27180520
AACAACGCCAGAA[TGTAGT-GTGGT--GGGAAAGC-insert-AGGGAGCGCGCATCTGGCTACA]CTAGGGACATGC
chr18:17015068-17018376
TTAGGAAAGCGAA[TGTAGC-GAGGT--GGGAAAGC-insert-AGGGAGCGCGCATCTGGCTACA]CATACTTGAGGC
```

**Similarities extend beyond 1.2-1.4 kb (More intact TE's)**

```
             [----------LTR--------->------TE------<------LTR------------]
chr14:3028070-3045206
TCAGCAATGCACA[TGTAGC-GAGGT--GGGAAAGC-longer_insert-AGGGAGCGCGCATCTGGCTACA]CACACATTCAGC
>chr23:8119468-8136406
ATTTTAATCCATA[TGTAGC-GAGGT--GGGAAAGC-longer_insert-AGGGAGCGCGCATCTGGCTACA]CATATTAAATGT
>chr20:27203891-27221089
CCCAATATGCTAG[TGTAGC-GTGGT--GGGAAAGC-longer_insert-AGGGAGCGCGCATCTGGCTACA]CTAGGGGACATG
>chr14:1845910-1863188
CATGGAAACATAA[TGTAGC-GTGGT--GGGAAAGC-longer_insert-AGGGAGCGCGCATCTGGCTACA]ATAACATTTTAT
>chr10:13237479-13254762
AGCACAATGTCAG[TGTAGC-GAGGT----------longer_insert-GTTGCCAACGCCACCTGGGGAATTTCA]TCAGCTTC
```
