## Supplementary Figures for "Heterochiasmy facilitated the establishment of *gsdf* as a novel sex determining gene in Atlantic halibut"

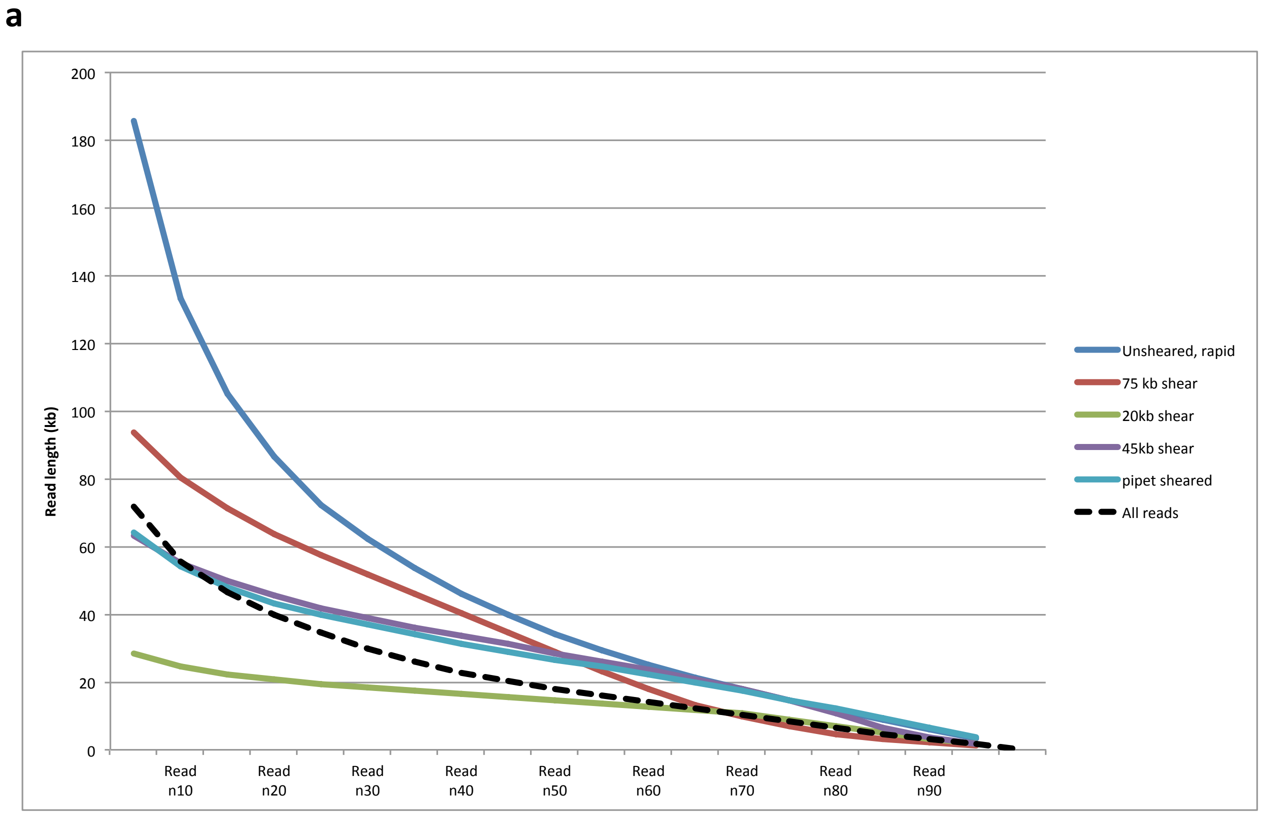


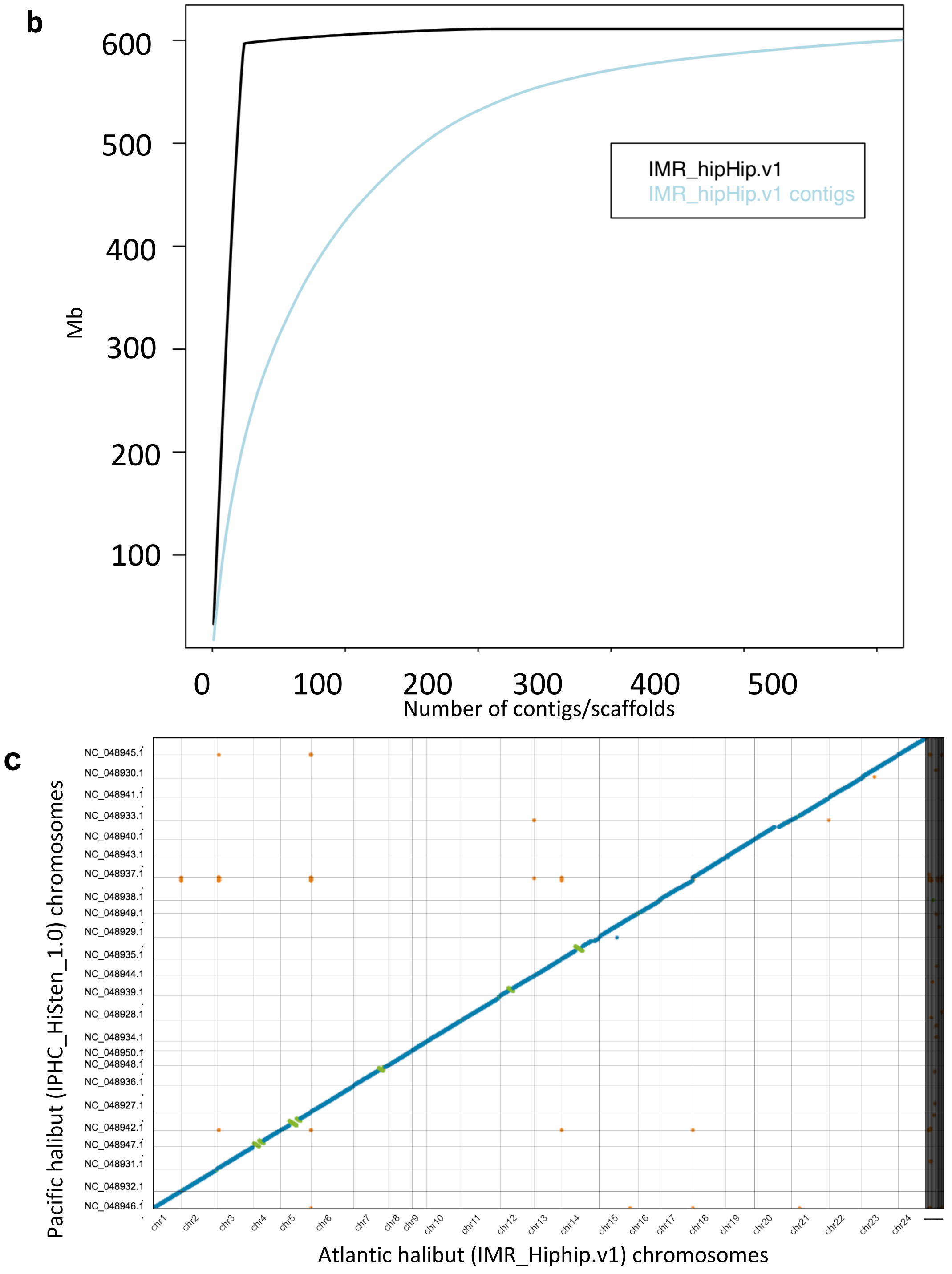


**Supplementary Fig. 1: a** Read length distribution of nanopore reads used to generate the IMR_Hiphip.v1 assembly. **b** Assembly continuities for the initial ONT contig assembly and the scaffolded IMR_Hiphip.v1 assembly. **c** Dot plot of genome to genome alignment between the IMR_hiphip.v1 assembly and the Pacific halibut reference genome assembly (IPHC_HiSten_1.0). Five chromosomes show larger inversions which are visible as green alignments perpendicular to the linear alignments (blue line) in the plot. A high degree of synteny was observed.

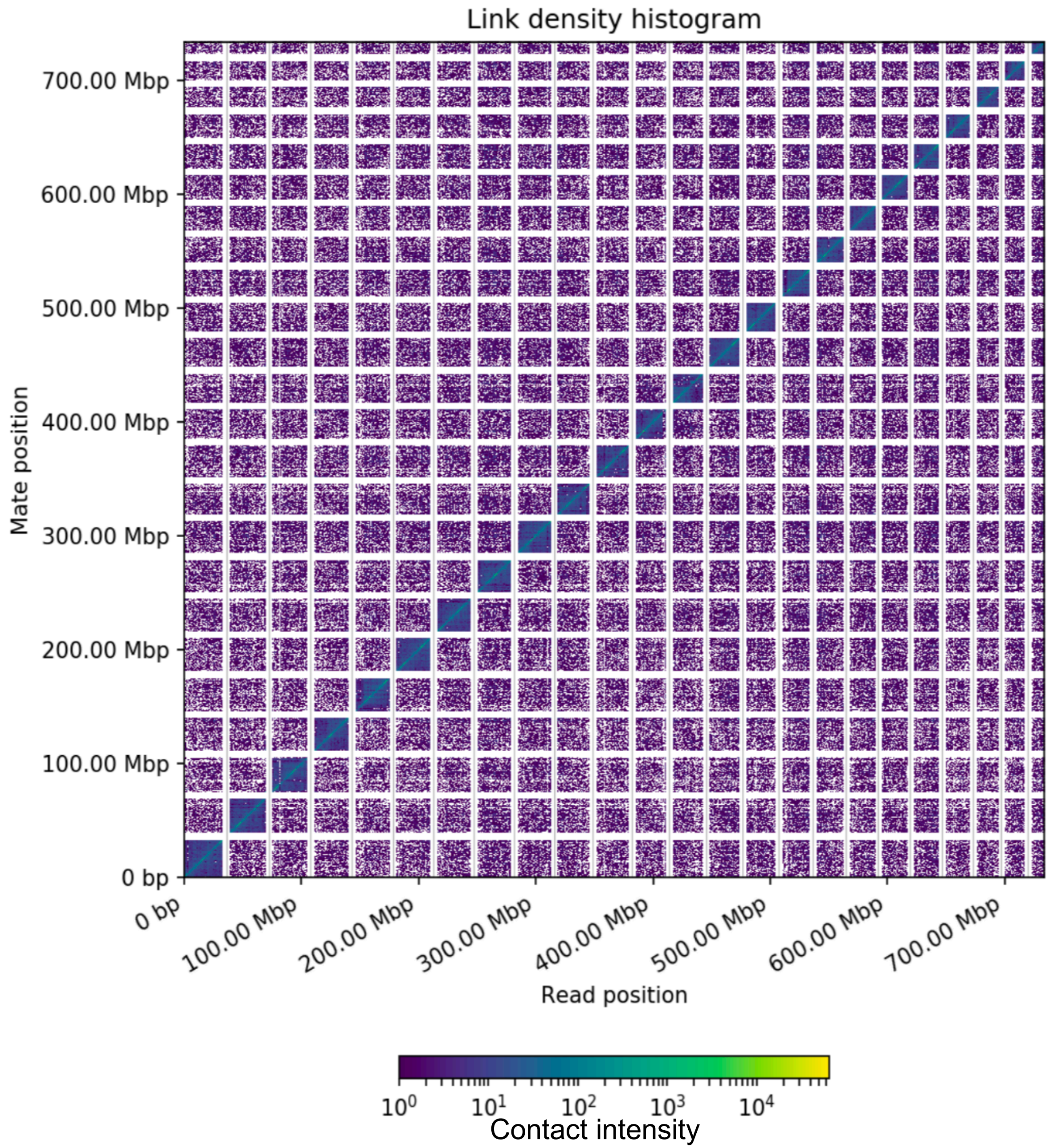


**Supplementary Fig. 2:** HiC contact matrix after scaffolding the Oxford Nanopore contig assembly using HiRise. Scaffolding resulted in 24 major scaffolds, in agreement with the karyotype of Atlantic halibut.

**Supplementary Fig. 3:**
Principal component analysis of the RNA-seq data reveals clustering by maturation stage rather than genetic sex. Dots indicate coordinates of individual RNAseq samples on PC1 and PC2. Sample sames indicate their ages in days post fertilization (dpf). Box colors indicate the genetic sex assignment of samples.

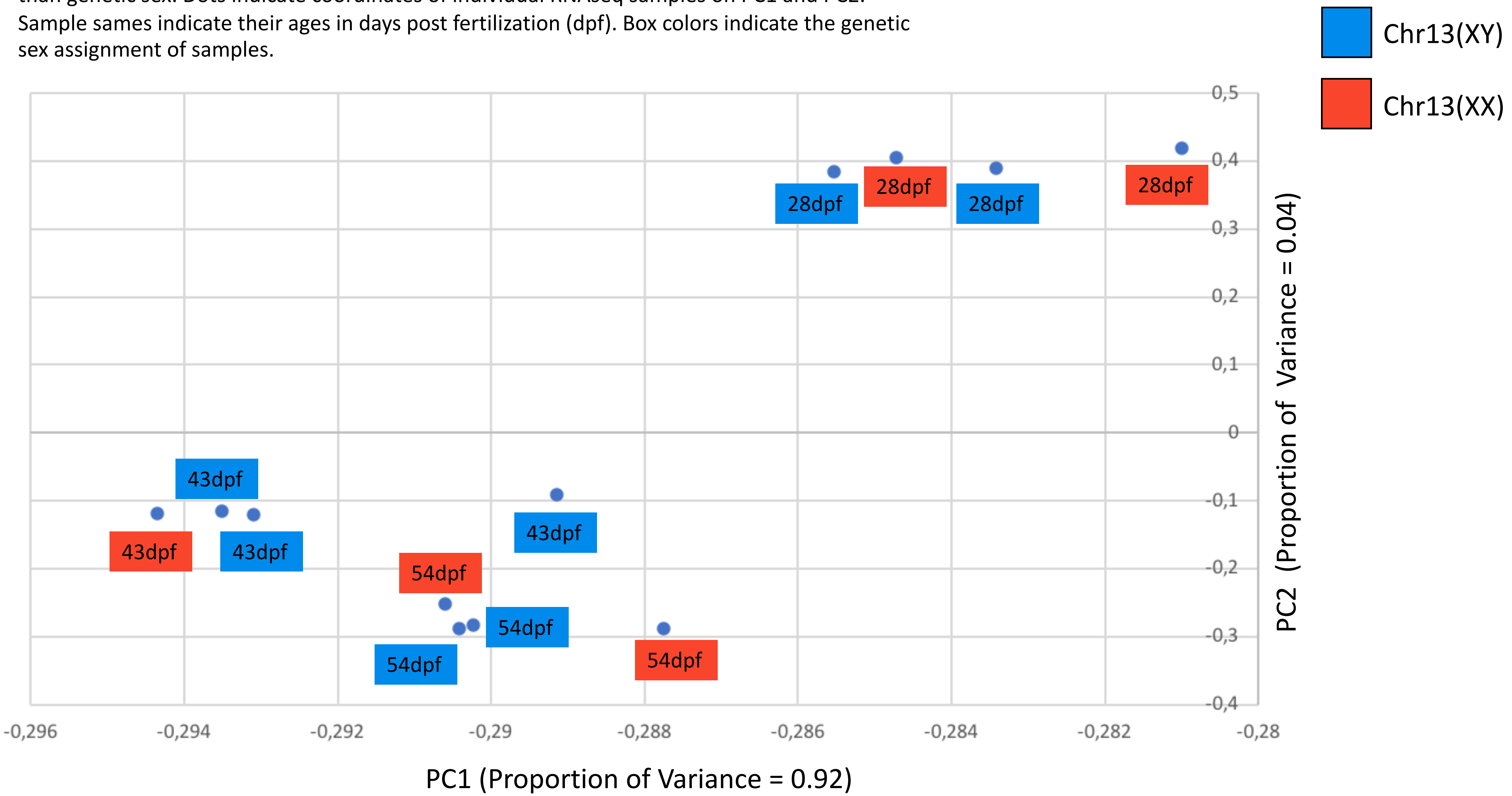

**Supplementary Fig. 4:**
**a** RNA-seq data samples clustered for 383 SNPs on chr13 previously found to be fixed in female DNA pool-seq while being variable in the male DNA pool-seq. We retained only positions were both alleles were observed in the RNA-seq data. Coordinates of genes along the Hiphip.v1 assembly are shown to the right. The samples without (NEG) *gsdf* expression in the RNA-seq and the samples with *gsdf* expression (POS) are in separate clusters. It was concluded that the individuals labeled XX and XY were female and male samples, respectively. **b** 65 SNPs fixed in the RNA-seq NEG samples but with both alleles observed in the RNA-seq POS samples along chr13. The nucleotide position (n) of the SNPs are shown to the right of the Figure. Made in https://software.broadinstitute.org/morpheus/

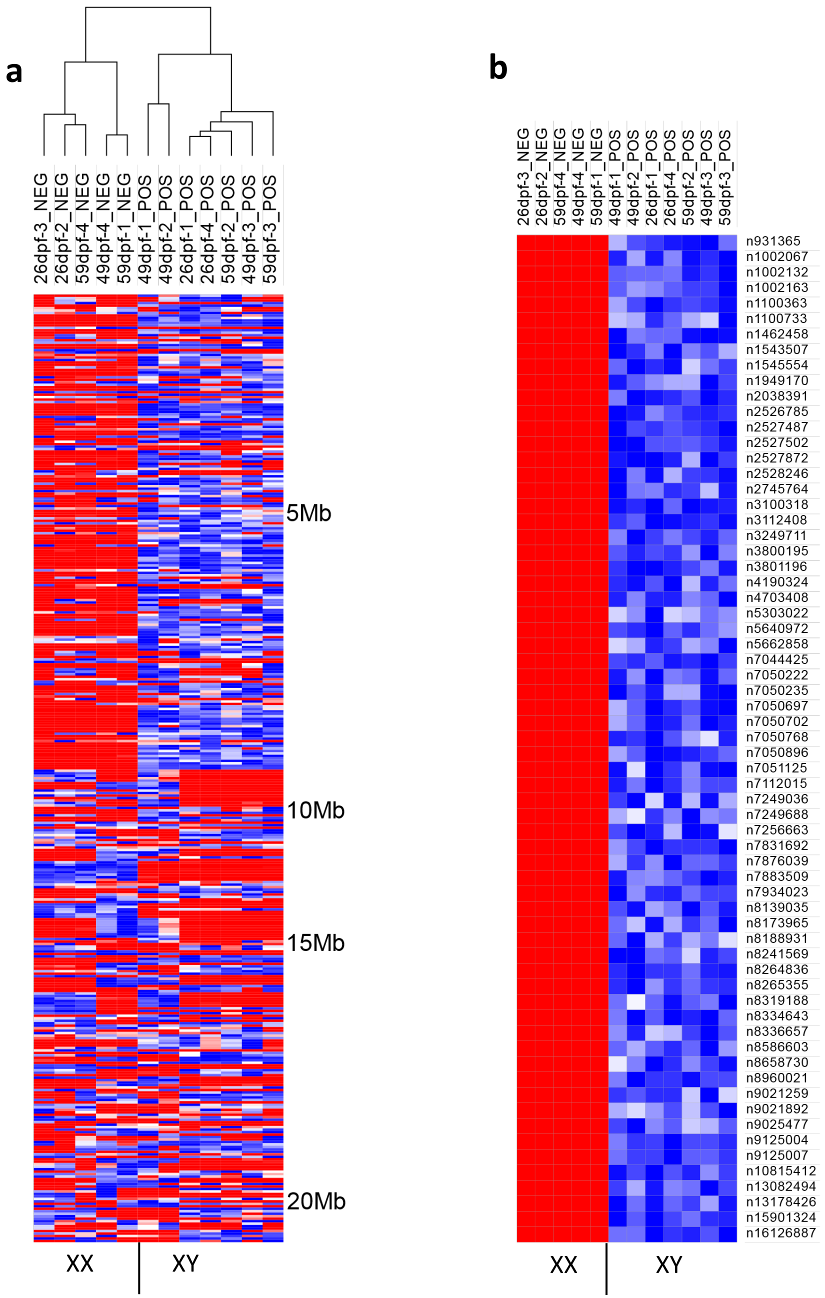

**Supplementary Fig.5: Genetic sex assignment by PCR.**
Sex differentiating PCR using sex specific primers (Table 1 in the manuscript) on cDNA from Atlantic halibut embryos at different developmental stages. Control samples were genomic DNA from individuals with known sex. Genetic sex of the samples was determined using a PCR based assay designed to differentiate genetic female and males. The forward primer was common for chr13X and chr13Y, while the reverse primers differed for two nucleotides in the 3'UTR of brx on chr13:9125004-9125007. For each individual sample (marked with a number) both female specific primers (brx_fwd + brx_femalerev; left well) and male specific primers (brx_fw + brx_malerev; right well) were run. In the case of a female sample, one band is detected in the left well while no band is detected in the right well. In the case of a male sample, two bands are detected. Genetic sex could not be determined for the 1, 8 and 24 hpf samples based on cDNA since this is prior to zygotic transcription of brx and the amount of gDNA extracted from these early stages were insufficient for reliable PCR.

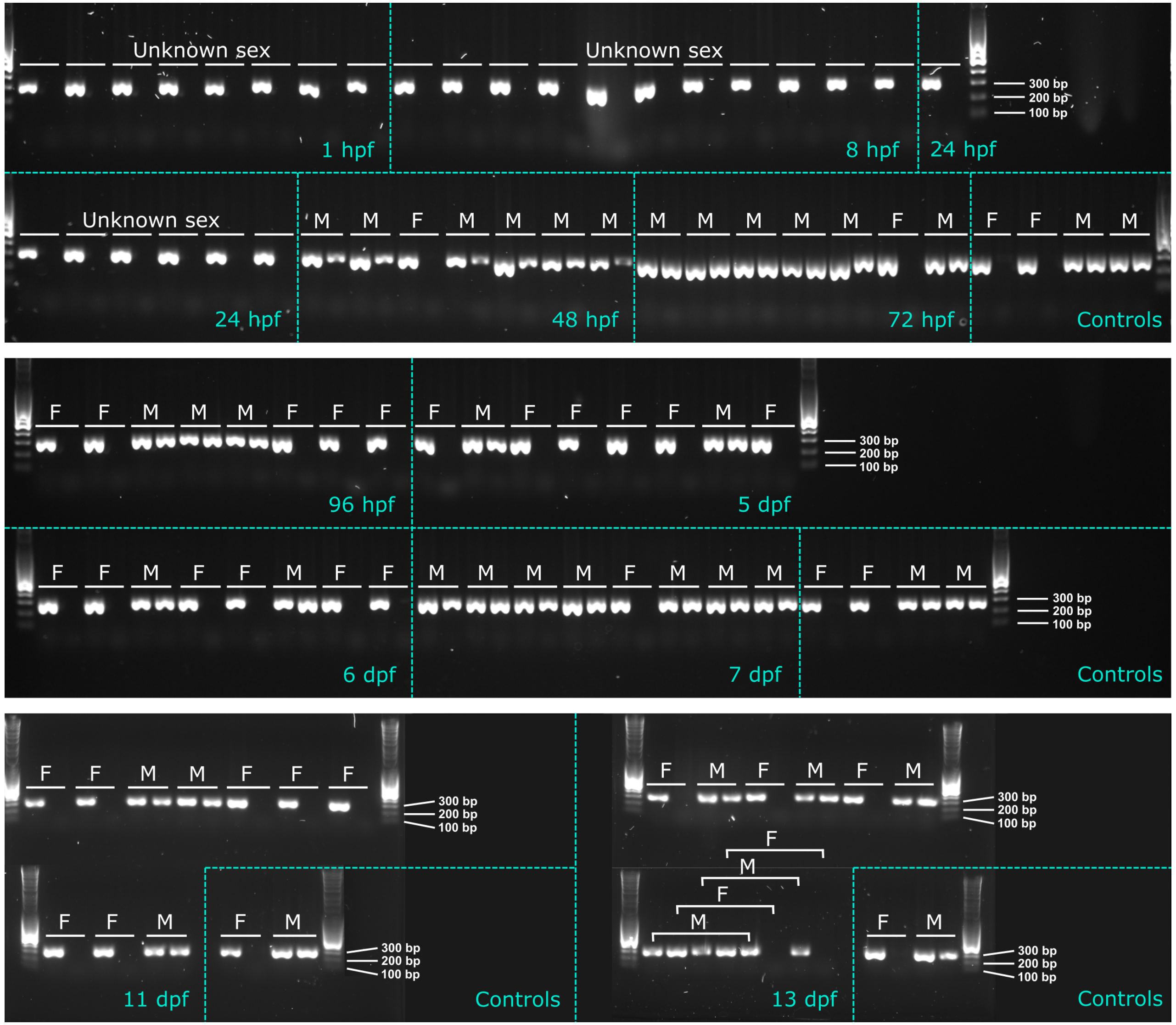

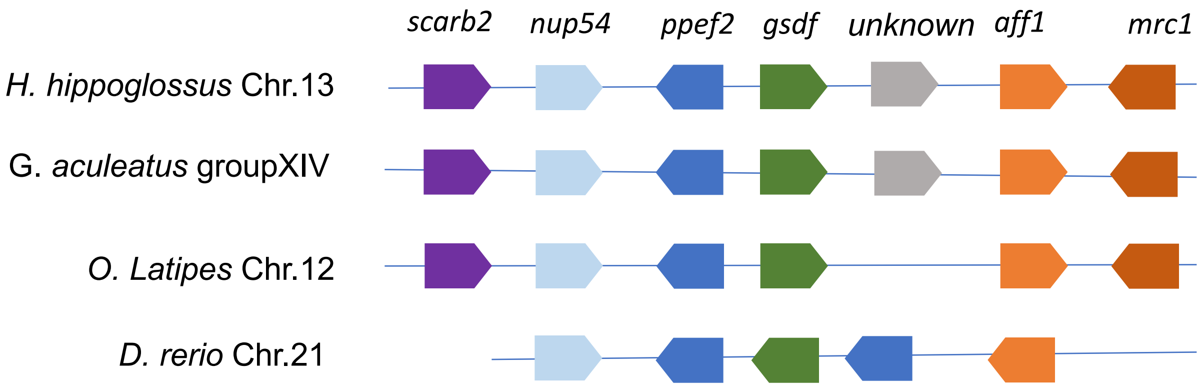


**Supplementary Fig. 6:** Synteny of chromosomal regions associated with *gsdf.* The diagram is not to scale for the lengths of each gene nor the distance between genes. The data was retrieved from http://www.ncbi.nlm.nih.gov and http://www.ensembl.org.

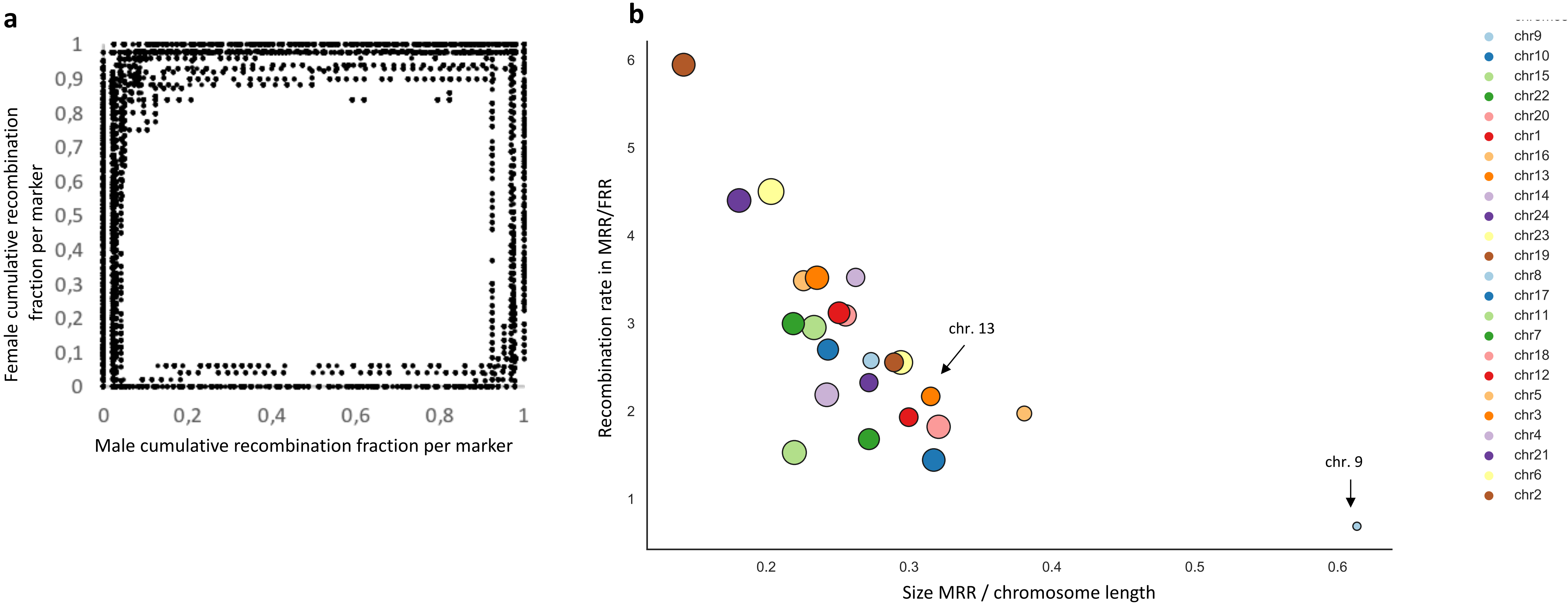


**Supplementary Fig. 7: a** XY-scatter plot of the cumulative genetic distances of chromosomally anchored linkage map markers in males and females reveals a large extent of heterochiasmy. **b** MRRs are smaller but have higher recombination rate. Differences in effective recombination rate (cM/Mb) between the Male- and Female Restricted Recombination regions (MRRs/FRRs) as a function of the fraction of chromosomal size annotated as MRR. Shown on the X-axis is MRR size divided by chromosome length. Shown on the Y-axis is the recombination rate (cM/Mb) for each MRR divided by the rate for the corresponding FRR. Circle sizes are proportional to chromosome lengths. The Atlantic halibut sex chromosome (chr13) is indicated by an arrow. Chr9, the smallest chromosome, is the only chromosome having a larger MRR than FRR as well as a higher female effective recombination rate

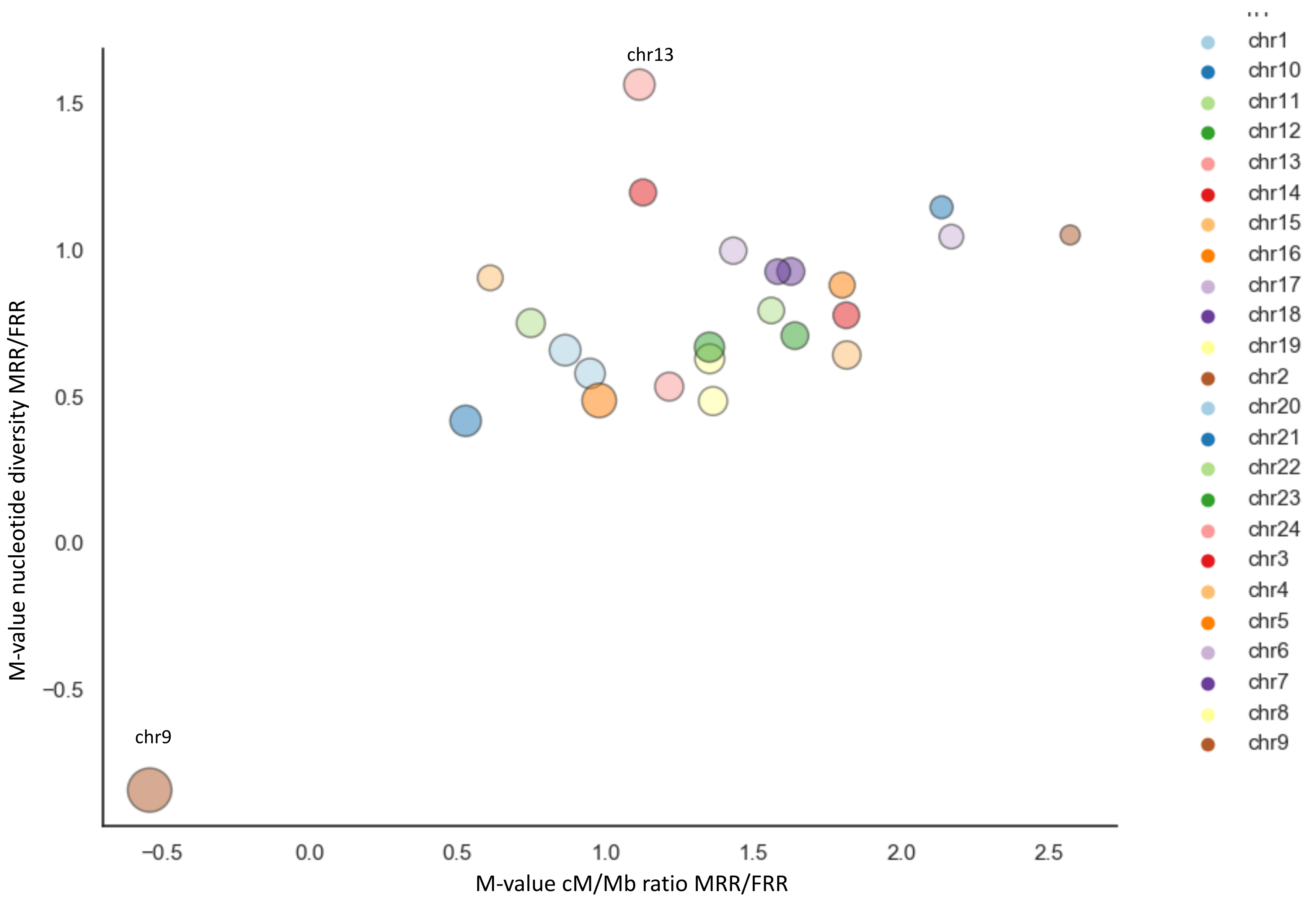


**Supplementary Fig. 8:**
The relationship between nucleotide diversity and recombination rate for MRRs and FRRs. On thex-axis: the log2 fold change (M-value) for recombination rate observed in MRRs and FRRs. On the y-axis: the log2 fold change (M-value) for nucleotide diversity in MRRs and FRRs. Circle colors indicate the chromosome and circle sizes are proportional to the relative size of the MRR on each chromosome (size MRR/ chr size). Chr13 (the sex chromosome) shows the largest difference of all chromosomes for MRR/FRR nucleotide diversity M-value. The FRR of chr13 is the X/Y chromosome.

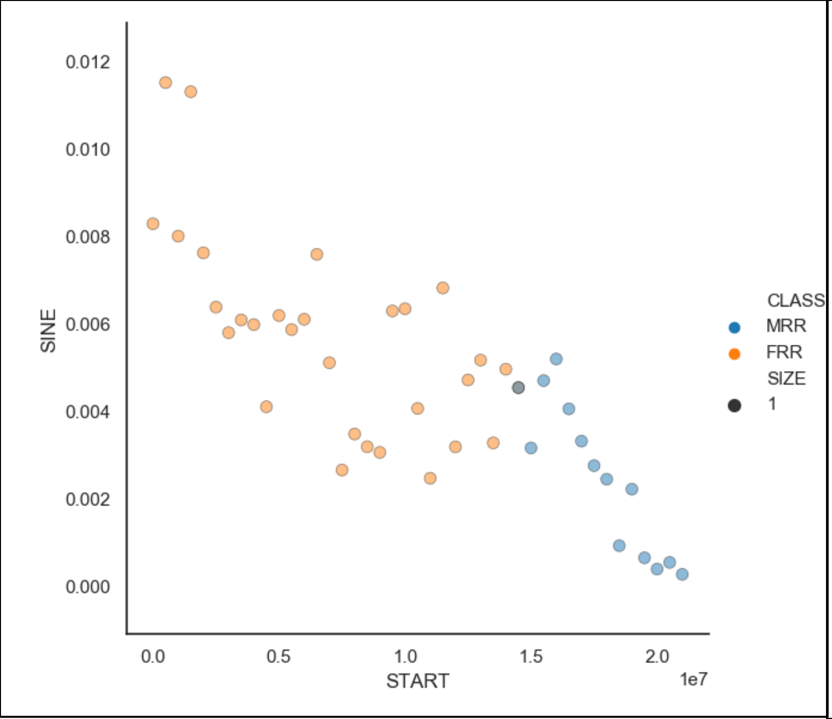


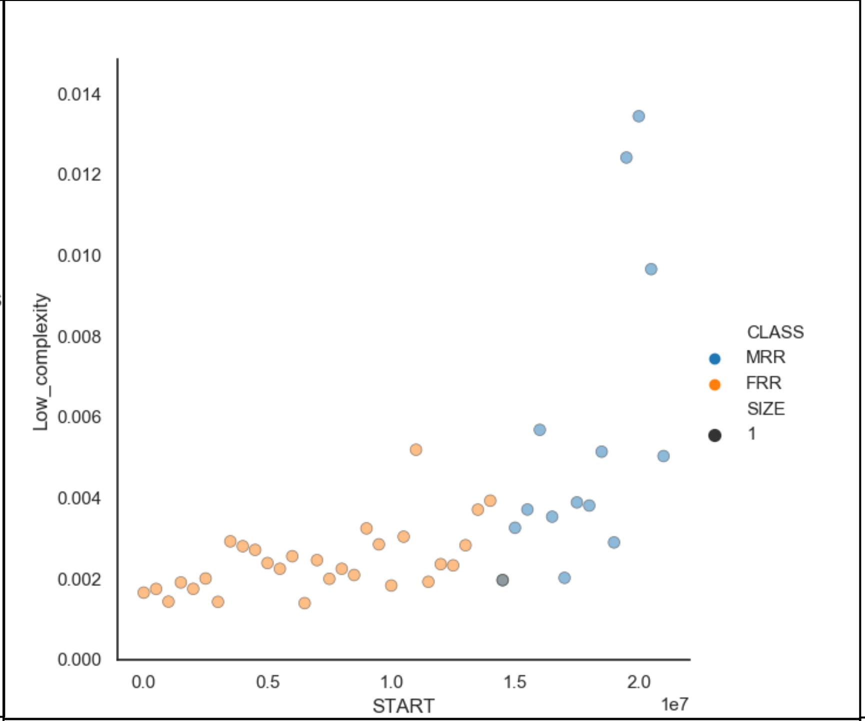


**Supplementary Fig. 9:**
Density of RepeatMasker classifications along the Atlantic halibut sex chromosome, chr13.
X-axes show coordinates along the chromosome ($10^7$ bp). Y-axes show the proportion of nucleotides in 500 kb windows overlapping an element belonging to each specific repeat superclass. Repeat superclass is indicated to the left of each subplot. Circle colors indicate whether a 500 kb window is classified as a MRR or FRR

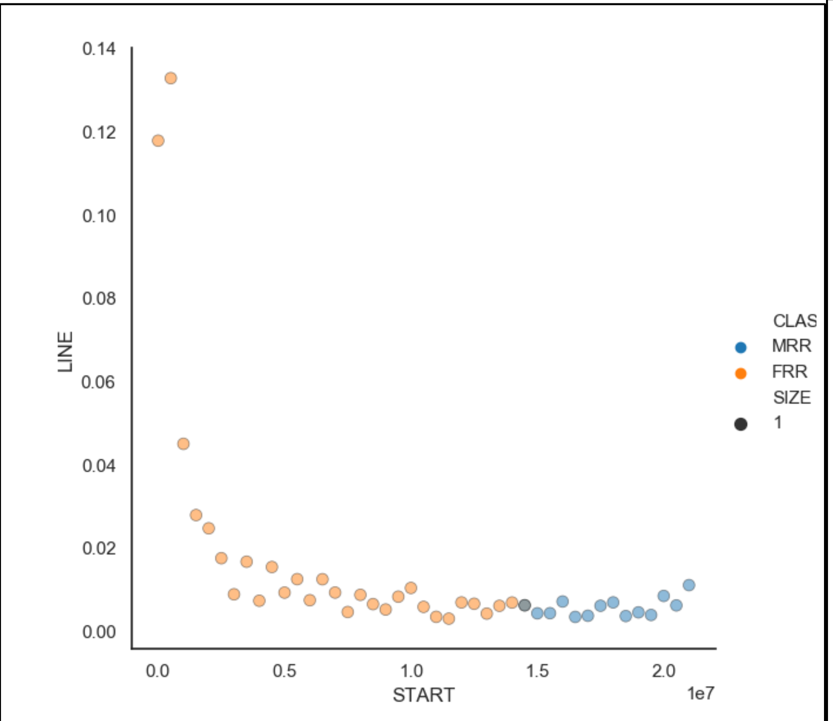


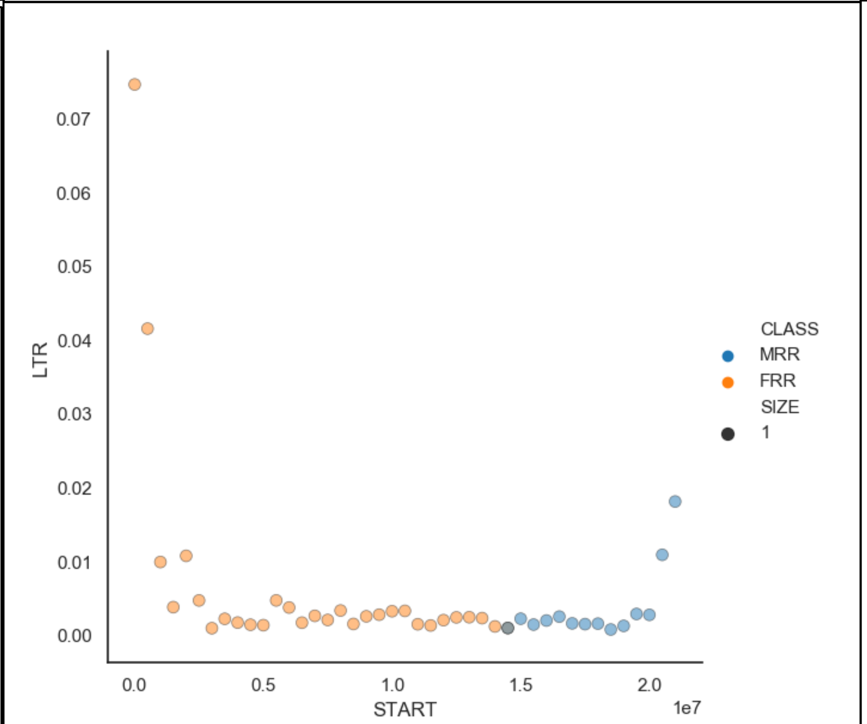


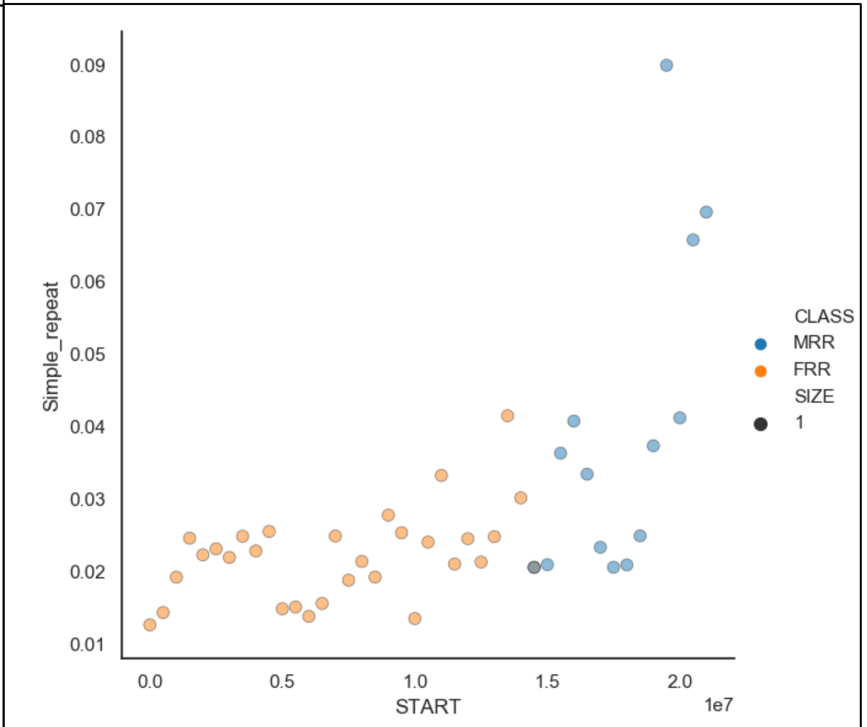

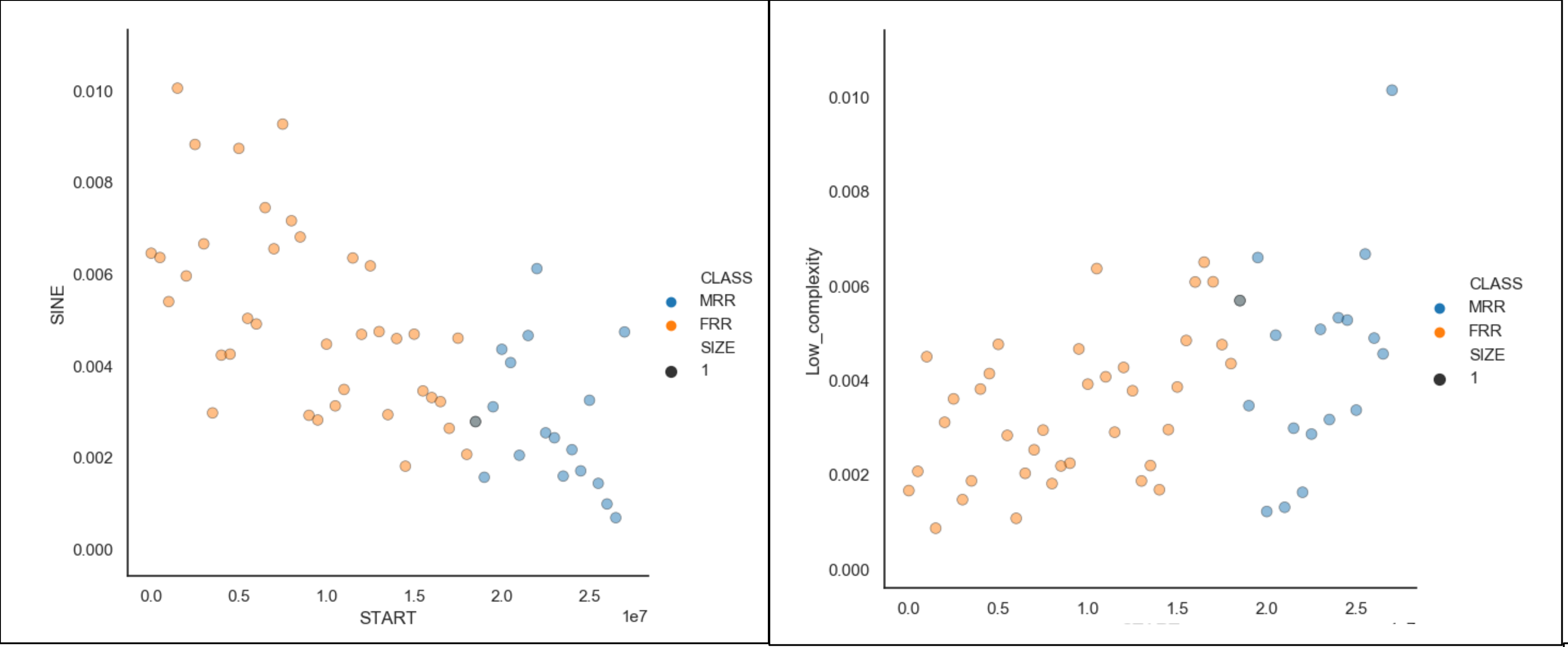


**Supplementary Fig. 10:**
Density of RepeatMasker classifications along Atlantic halibut autosome chr10.
X-axes show coordinates along the chromosome ($10^7$ bp). Y-axes show the proportion of nucleotides in 500 kb windows overlapping an element belonging to each specific repeat superclass. Repeat superclass is indicated to the left of each subplot. Circle colors indicate whether a 500 kb window is classified as a MRR or FRR.

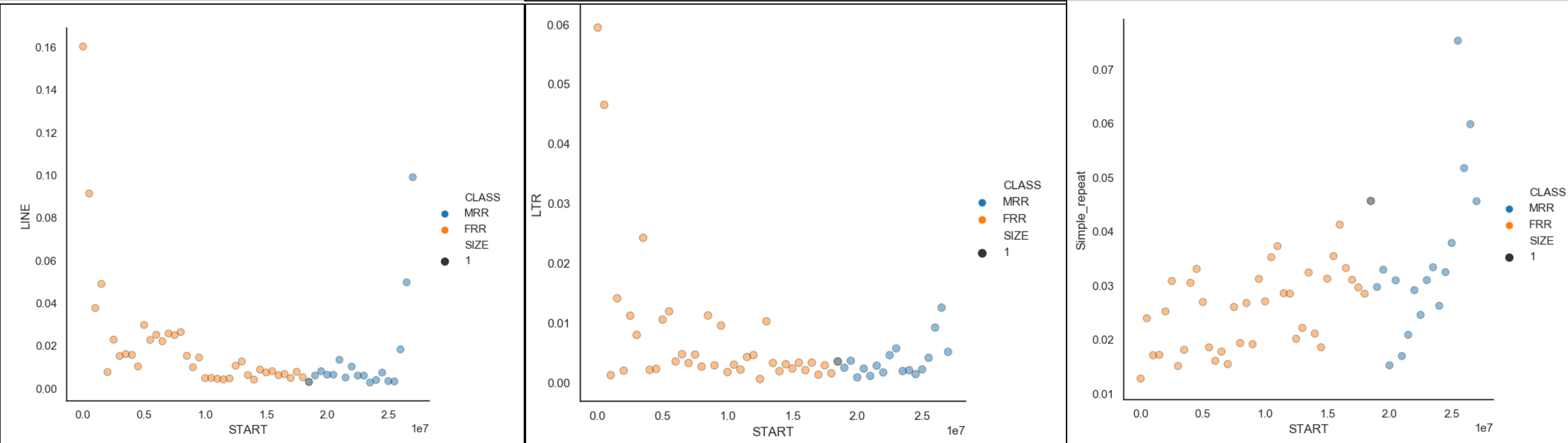

**Supplementary Fig. 11**

HiC contact maps (all chromosomes and chr13 speparately) of the male (XY) Atlantic halibut used to generate the genome assembly. No apparent large-scale inversions or translocations were observed (would appear as off-diagonal contacts)

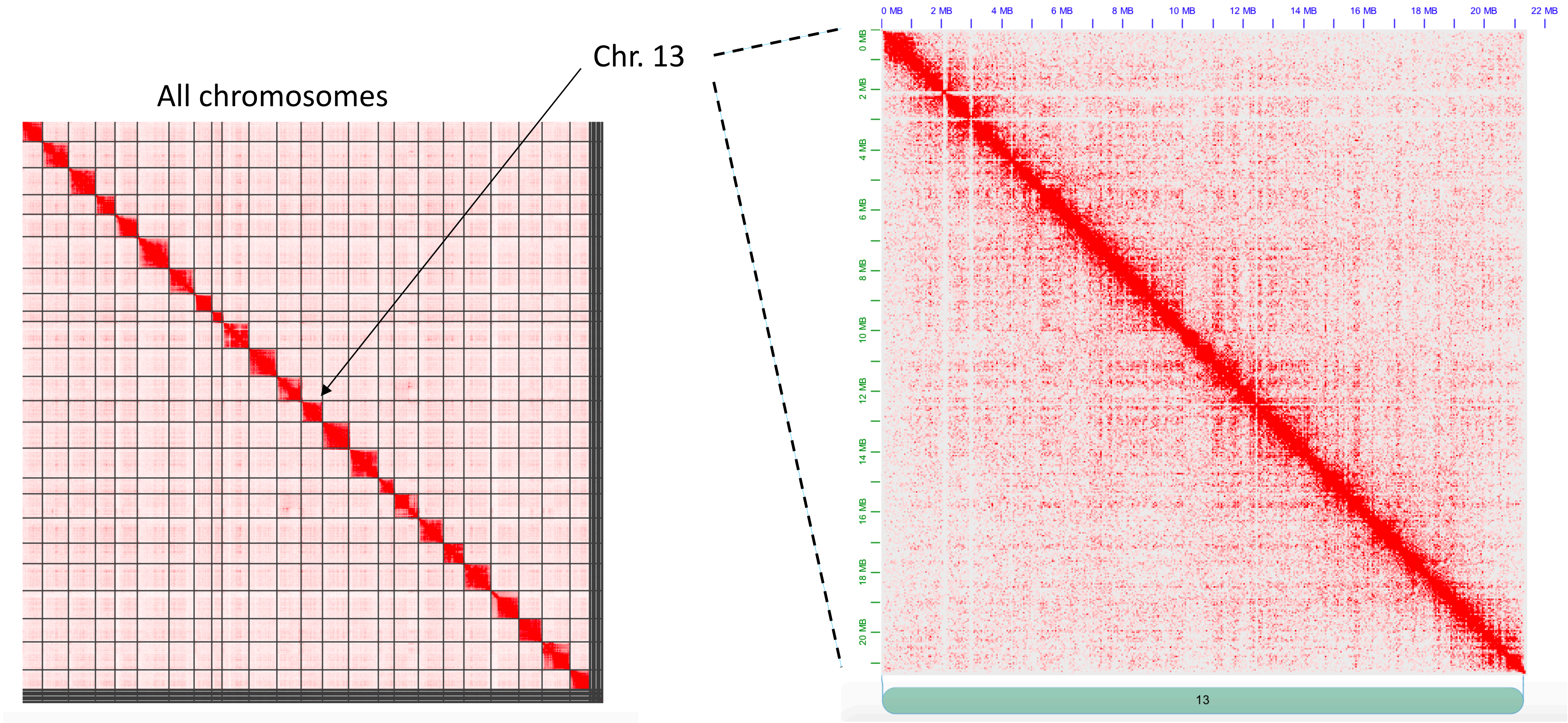

**Supplementary Fig. 12:**

*Chr. Y carries a transposon-derived LTR upstream of gsdf*

Male- and female haplotypes differ for a 1.2 kb transposable element derived segment 2 kb upstream of *gsdf* transcription start site of. Males carry the insertion allele which contains a lowly methylated CpG-island inferred to act as a derived promoter of *gsdf* uniquely in males. Shown are alignments of male- and female poolseq data, nanopore reads and supernova pseudo-haplotype assemblies from the single male used to create the IMR_Hiphip.v1 assembly. A heterozygote 1.2 kb insertion is evident in nanopore read alignments as well as in the linked-read haplotype alignments and chr13Y carries the insertion allele. The insertion is less evident in the male short-read data due to poor mappability in the in/del region. Male pool short reads carrying chr13Y haplotype variant tags have fever red colored mates supporting the deletion allele (short read alignments are sorted by the SNP allele in the center red/blue box, blue boxes= chrY-allele, red boxes=chrX-allele). *gsdf* gene model in blue shows the extent of *gsdf* expression detected in Atlantic halibut adult testis and adult ovary and testis in Pacific halibut. The two *gsdf* gene models in red indicate isoforms detected only in males from three separate developmental stages.

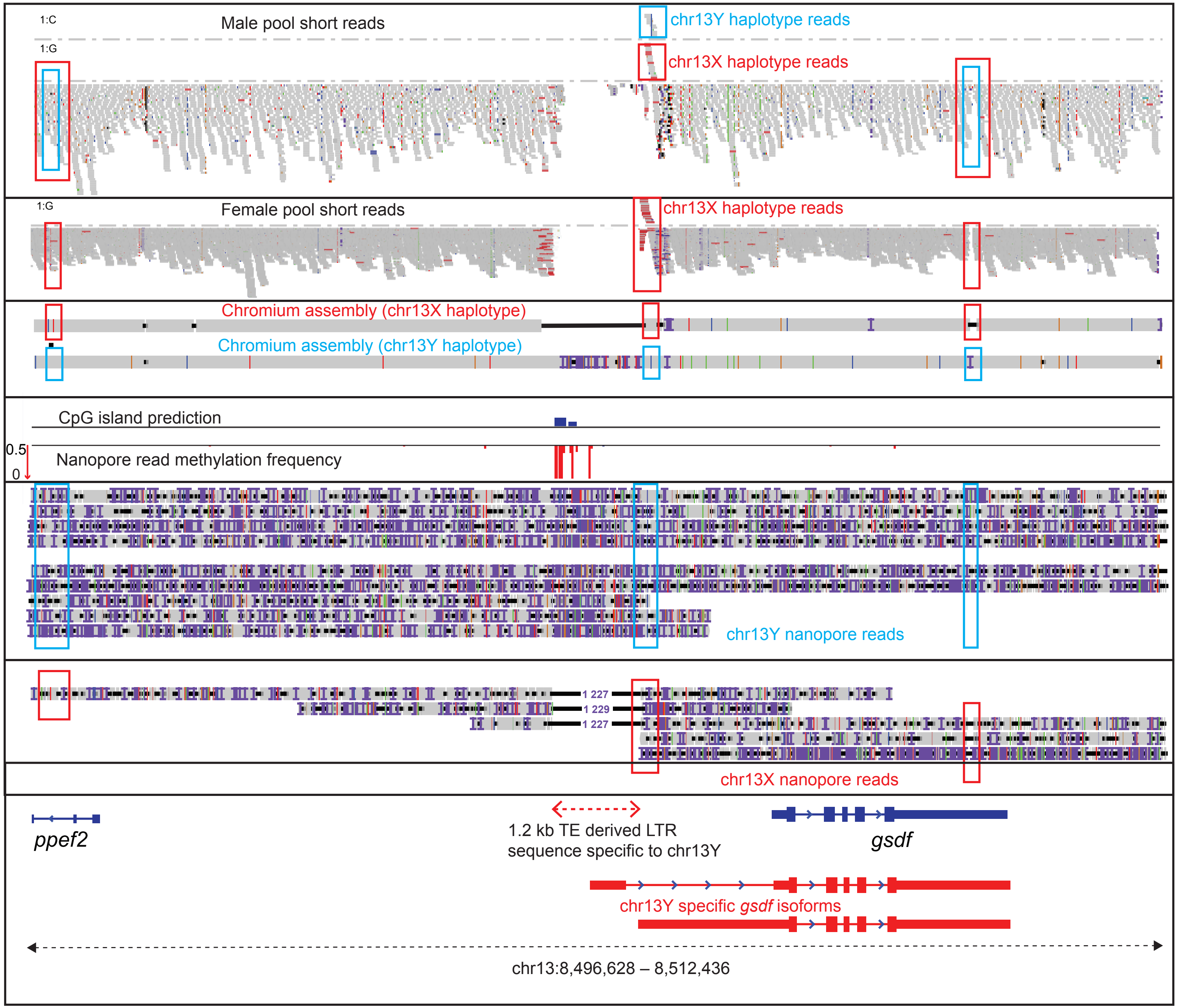

### Supplementary Fig. 13

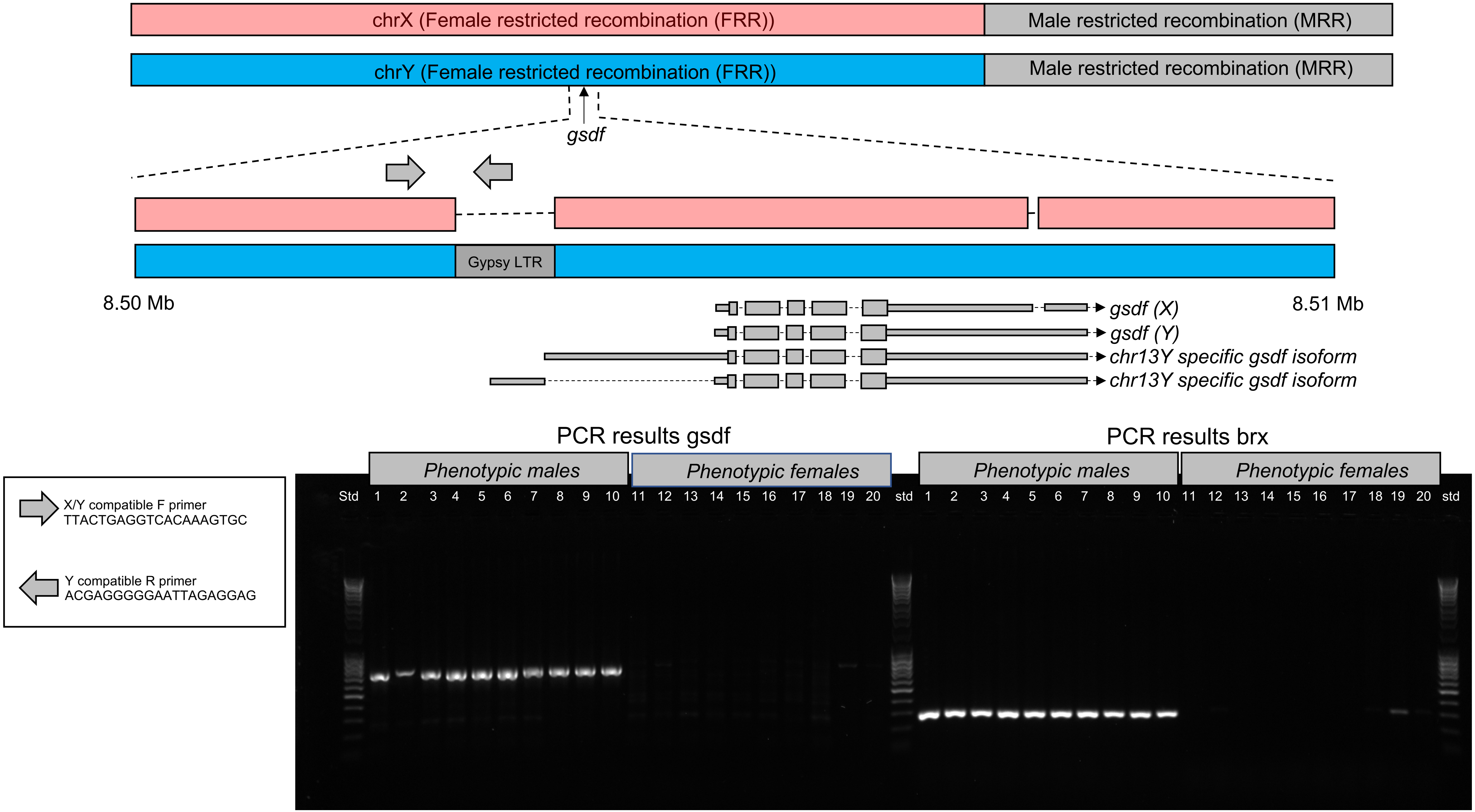

**Supplementary Fig. 13:** Schematic representation of *gsdf* and *brx* PCR results. The forward *gsdf* primer is 481 nt upstream of the 1.2 kb chrY Gypsy-LTR insertion. The reverse primer is 251 nt into the LTR. The *brx* primers are described in Methods. The resulting PCR products are shown in an agarose gel. The same 20 individuals are shown for both primer pairs. Individuals 1-10 are phenotypic males and 11-20 are phenotypic females. PCR results show that *brx* classification and presence of Gypsy-LTR insertion are in agreement and distinguish all 10 males from females.

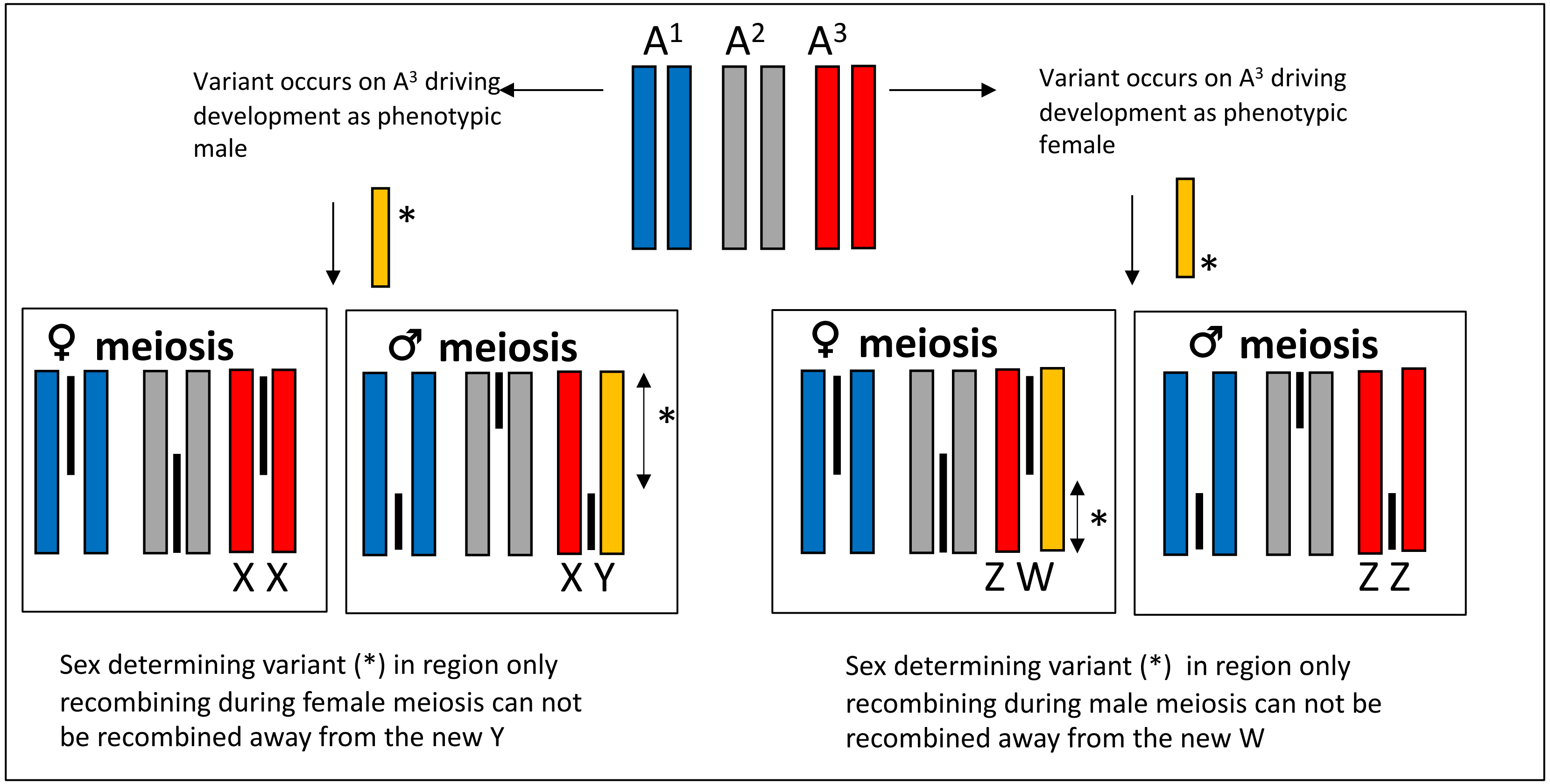


**Supplementary Fig. 14:** Theoretical evolution of sex chromosomes in a heterochiasmy setting. On top, three fictive autosome pairs ($A^1$-$A^3$) are shown. One of these aquires genetic material governing development of carriers into phenotypic males (left arrow) or phenotypic females (right arrow), thereby creating a novel sex chromosome. Thick black lines in between chromatids indicate the chromosomal interval undergoing meiotic recombination in each sex. The new sex determining gene (*) must occur in a region incapable of meiotic recombination in the heterogametic sex, or else, it does not become isolated from recombining with it's former autosomal pair during meiosis. Arrows within boxes next to the sex determining gene location indicate the interval where a new sex determining gene could occur in the two respective systems, (XY or ZW).

**Supplementary Fig. 15:** Reference gene for qPCR. To find a suitable (stable) internal reference gene for the normalization of the qPCR data, we selected a few candidates from the RNAseq dataset with a stable number of reads throughout the developmental stages available. **a** Expression of the selected genes *ints9*, *JKAMP* and *gtf3c6* in Atlantic halibut  embryos and larvae, shown as number of reads. Data are shown as mean with SEM. N=4. **b** The selected genes *ints9*, *JKAMP* and *gtf3c6* were selected for testing with qPCR on the actual samples to be analyzed for *gsdf* expression. Expression of in Atlantic halibut embryos and larvae, shown as cycle threshold (CT)-values. Data are shown as mean with SEM. N=4-9. *gtf3c6* had the most stable expression throughout the developmental stages, and was therefore chosen as the internal reference gene in this study.

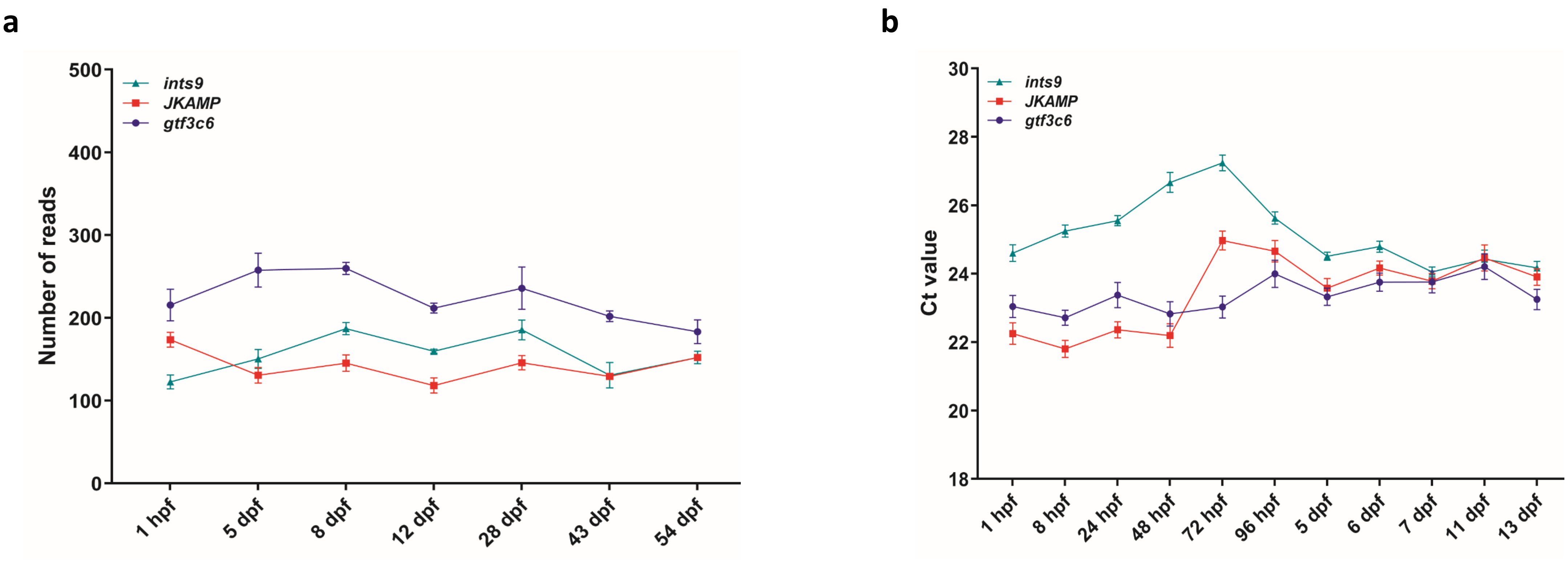
